# Transcriptional analysis of immune modulatory genes in melanoma treated with PD-1 blockade

**DOI:** 10.1101/2020.12.20.397000

**Authors:** Hyojin Song, Sungyoung Lee, Se-Hoon Lee, Miso Kim, Sang Yup Lee, Sung-Soo Yoon, Hongseok Yun, Youngil Koh

**Affiliations:** Center for Precision Medicine, Seoul National University Hospital, Seoul, Republic of Korea, 03080; Cancer Research Institute, Seoul National University College of Medicine, Seoul, Republic of Korea, 03080; Center for Medical Innovation, Seoul National University Hospital, Seoul, Republic of Korea, 03080; Division of Hematology-Oncology, Department of Medicine, Samsung Medical Center, Sungkyunkwan University School of Medicine, Seoul, Republic of Korea, 06351; Department of Health Sciences and Technology, Samsung Advanced Institute of Health Science and Technology, Sungkyunkwan University, Seoul, Republic of Korea, 06351; Department of Internal Medicine, Seoul National University Hospital, Seoul, Republic of Korea, 03080; Department of Chemical * Biomolecular Engineering, Korea Advanced Institute of Science and Technology, Daejeon, Republic of Korea, 34141

**Keywords:** Immune checkpoint modulator, Transcriptional signatures, Regression-based modeling, Feature selection, Immune status measure, Immune temperature, Melanoma

## Abstract

We aimed to characterize immunological features of melanoma patients treated with PD-1 blockade using tumor transcriptomic datasets. Response-dependent and response-independent predictors based on biological knowledge were investigated. Domain knowledge-driven regression-based analysis identified *CEACAM1, CD40, B7-H3*, and *CD112* as key genes that determine the melanoma immune status. We devised the transcriptional deviance score (*TDS*) representing the individual sample-wise contribution to the immune network. The *TDS* not only showed good predictive power for immune checkpoint inhibitor (ICI) responses but also suggested specific gene interactions that determine ICI responses. Dynamic *TDS* changes following ICI treatment were related to long survival, indicating immune network modulation by ICIs occurred in responders. A predictive model incorporating *B7-H3* and *CEACAM1* expression, mutational status, clinical features, and the *TDS* showed excellent performance for ICI response. Thus, our approaches suggest a novel measure for the tumor immune temperature and provide insight into melanoma immunobiology.

**Highlights:**

- We applied outcome-independent and outcome-dependent methods to investigate melanoma immunobiology.
- *CEACAM1, CD40, B7-H3*, and *CD112* expression levels are key determinants of immune status.
- We devised a *TDS* that could measure tumor immune network status at the individual level.
- Incorporating regression and correlation approaches greatly improves predictive power.

## Introduction

Immune checkpoints directly or indirectly regulate the ability of T cells to recognize and attack cancer cells. The advent of immune checkpoint modulators significantly improved the survival of cancer patients (Ribas, 2012; Ribas et al., 2003). Specifically, immune checkpoint inhibitors (ICIs) targeting the PD-1/PD-L1 or CTLA-4 axis are widely used for the treatment of several cancer subtypes. However, the response rate of pembrolizumab is approximately 20% in non-small cell lung cancer (NSCLC) (Garon et al., 2015) and 30% in advanced melanoma (Robert et al., 2015). Given the limited response rate in cancers, understanding the biological mechanism of the immune checkpoint modulator response is fundamental to optimize the use of ICIs in cancer patients.

Two recent studies have reported on response patterns to anti-PD-1 therapy in melanoma. Hugo et al. (Hugo et al., 2016) discovered influential factors on the basis of somatic mutations and transcriptional signatures. They observed an association between higher mutational load and improved survival in metastatic melanoma and introduced the innate anti-PD-1 resistance (IPRES) signature. Riaz et al. (Riaz et al., 2017) demonstrated evolutionary dynamics in the tumor microenvironment modulated by anti-PD-1 therapy (nivolumab) through integrated analysis of whole-exome, transcriptome, and T cell receptor sequencing datasets. They identified reduced mutational and neoantigen loads in anti-PD-1 responders and further observed increased immune cell subsets, transcriptional network activation, and upregulated immune checkpoint expression in longitudinal transcriptomic analyses. These studies provide comprehensive information on the genomic and transcriptomic patterns found in melanoma patients receiving PD-1-axis blockade.

On the other hand, predictive measures for outcomes after PD-1/PD-L1-targeted treatment remain unsatisfactory. In fact, melanoma is considered a cancer type with highly variable immunogenicity (Roh et al., 2017; Twitty et al., 2020). So far, three predictive biomarkers for ICI in melanoma have been repeatedly suggested: PD-L1 expression on tumor cells, tumor mutational burden (TMB), and tumor-infiltrating lymphocytes (TILs) (Darvin et al., 2018; Kitano et al., 2018; Teng et al., 2015). First, positivity for PD-L1 expression at a threshold of 1% is reported to be associated with responsiveness and survival rates in solid cancers (Ancevski Hunter et al., 2018; Kim et al., 2018; Morrison et al., 2018; Sidaway, 2018). Despite the predictive roles of PD-L1 expression, the heterogeneity within tumors (McLaughlin et al., 2016) and local inflammation triggered by inhibitory treatment (Vilain et al., 2017) affect the expression levels of PD-L1. Hence, at this point, PD-L1 positivity must be further investigated to improve its predictive power for treatment response (Davis and Patel, 2019; Sorensen et al., 2016). Second, a high TMB, measured as the number of nonsynonymous mutations per megabase, may predict the survival but not the responses of melanoma patients treated with PD-1 inhibitors (Alexandrov et al., 2013; Hugo et al., 2016; Luke et al., 2017). Although Hugo et al. observed a higher median for the number of nonsynonymous SNVs in anti-PD-1 responders than in nonresponders (Hugo et al., 2016), this difference did not reach statistical significance. Thus, the use of TMB in melanoma needs statistical refinement and further adjustment (Keenan et al., 2019). Last, immune-suppressive cells, including TILs, regulate antitumor immune responses, and a correlation between increased numbers of TILs and the clinical benefits of ICIs has been observed (Geukes Foppen et al., 2015; Nishino et al., 2017; Tumeh et al., 2014). Despite their biological significance, immune infiltrates are very heterogeneous; therefore, there are limitations in the clinical measurement of TILs due to their heterogeneity. In fact, TIL counts fluctuate depending on the spatial location among tumor cells, even within the same tumor type (Fridman et al., 2012).

From an immunological perspective, cancers have been recently classified as either hot or cold tumors, which can be represented as the tumor immune temperature. Accordingly, recent studies have used diverse approaches to understand the tumor immune temperature. A recent study by Galon and Bruni et al. introduced 4 tumor classifications (hot, altered-excluded, altered-immunosuppressed, and cold) (Galon and Bruni, 2019) to indicate the immunological status of the surrounding microenvironment based on the concept of hot and cold tumors. Studies scoring immune temperatures, such as the tumor inflammation signature (TIS) (Danaher et al., 2018) and Immunoscore (Galon et al., 2016), are other recent efforts. However, the association between ICI responses and these conceptual scores has not yet been clearly established.

Considering these unmet needs related to our understanding of the immune mechanism underlying the ICI response in melanoma, we tried to dissect immune signatures in anti-PD-1-treated melanoma patients using transcriptomic datasets. Since our immune system has substantially dynamic aspects, we believed that RNA sequencing (RNA-seq) data would be a particularly useful way to represent the tumor immune status. Hence, we sought to devise a method to adequately interpret the immune features with regard to PD-1 blockade from RNA-seq datasets. Our study uses two approaches that employ regression- and correlation-based frameworks. These approaches utilize machine learning techniques combined with domain knowledge-based biological information. We assumed that dimension reduction would deduce the essential implications of the immune modulators that interact with each other by either forming networks or functioning independently. Hence, with the regression-based approach, we implemented a combination of feature selection and a machine learning analytic framework to reduce input variables (i.e., genes) for predictive modeling and extract transcriptomic determinants that contribute to gene expression patterns. With the correlation-based approach, we devised a new calculation method to measure the interactive status of immune checkpoint expression by quantitating individual contributions to the correlation. We assumed that the association between the expression levels of immune checkpoint modulators may act as a decisive index that reflects the mechanical concepts of hot and cold tumors.

To understand how the status of immune engagement in the tumor microenvironment is affected by PD-1 blockade, we developed a novel measure to quantify and classify the immune status and introduce transcriptomic determinants for PD-1 inhibition in melanoma. Our results may suggest a new point of view on anticancer treatment by enhancing immunotherapeutic strategies with the selection of druggable biomarkers.

## Results

### The established basal gene set covers 38 immune checkpoint modulators

We aimed to identify how immune checkpoint modulators and their interactions explain the effect of ICIs by using our analytic workflow (**Figure 1**). First, we established the basal gene set of 38 immune checkpoint modulatory genes (21 receptors, 16 ligands, and 1 receptor/ligand; **Table S1**) through in-depth literature curation (Andrews et al., 2019; Charoentong et al., 2017; Chretien et al., 2019; Galon and Bruni, 2019; Lines et al., 2014; Ott et al., 2013; Wang et al., 2014; Zarour, 2016). The gene set includes receptor-ligand pairs whose functions and mechanisms were determined in previous studies. Throughout the study, we conducted the entire analytical procedure (**Figure 1A**) based on the selected gene set. Specifically, in both the regression (**Figure 1B**) and correlation (**Figure 1C**) approaches, the transcriptomic data quantified in transcripts per million (TPM) were used to develop predictive models to understand the varied responses to PD-1 blockade. In particular, we employed both approaches to select predictors and conduct balanced model development in both response-dependent and response-independent frameworks.

**Figure 1.**
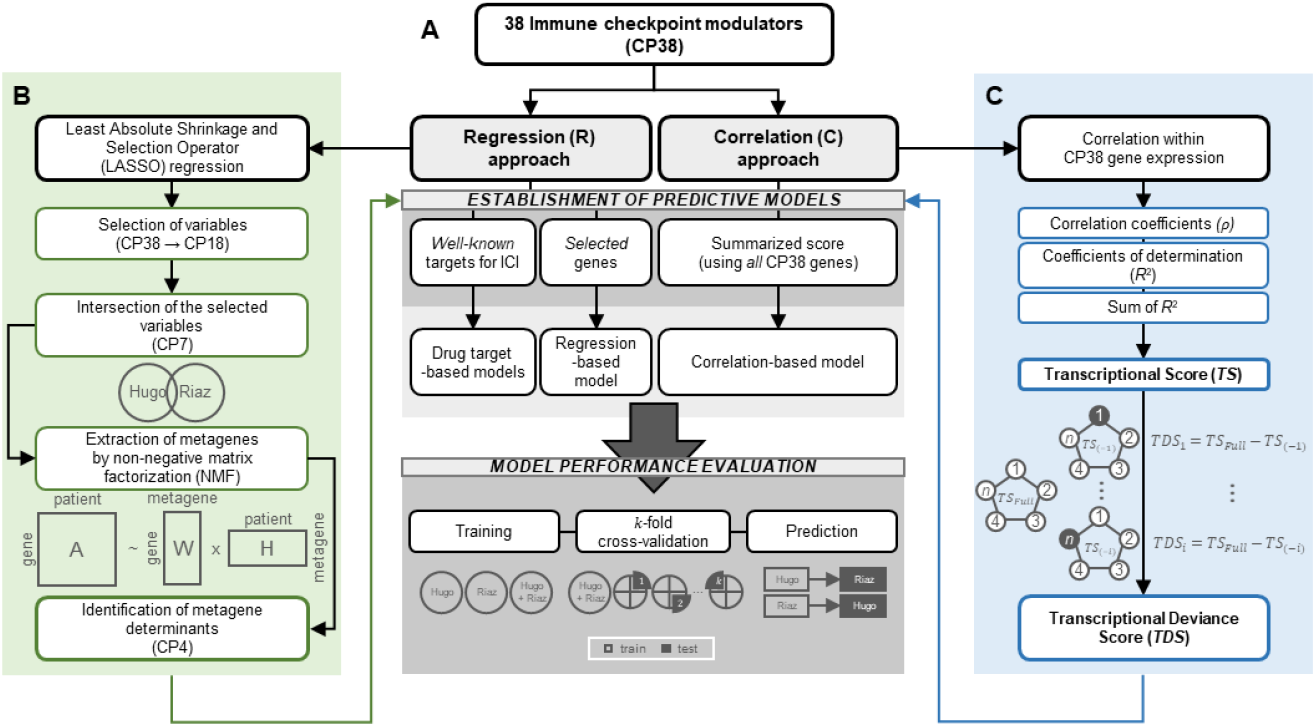
Workflow of the study. (A) The overall workflow of the study. The workflow briefly illustrates the two approaches of the regression- and correlation-based frameworks used in this study. (B) The detailed workflow of the regression-based approach including LASSO regression and the NMF method. From LASSO, 7 common variables (i.e., genes) were selected. Afterward, 4 metagene determinants were extracted from NMF based on the common genes. (C) The detailed workflow of the correlation-based approach including the conceptualization of *TS* and *TDS*. The *TDS* for the *i*^th^ individual is calculated by subtracting *TS*_*(-i)*_ from *TS*_*Full*_, where *TS*_*(-i)*_ represents the *TS* calculated without the *i*^th^ sample subtracted from the entire sample and *TS*_*Full*_ represents the *TS* calculated by using the entire sample.

### Feature selection using LASSO showed substantial agreement across datasets

In the regression approach, we focused on how the traditional regression-based approach explains the effect of PD-1 blockade. Principal component analysis revealed that both of our target melanoma datasets (see **Methods**) harbor a mild batch effect, and we were unable to completely correct those effects. The batch effect entailed distinctive dataset-specific patterns (**Figure S1**). Feature selection with least absolute shrinkage and selection operator (LASSO) regression of the two independent datasets yielded the 18 selected genes for each dataset, which is the optimal number to select representative features related to anti-PD-1 responses. We then compared the selected genes yielded from each dataset and identified 7 genes (*B7-H3, CD137, CD137L, CD112, CD40, CEACAM1*, and *KIR3DL1*) that were included in both datasets (**Figure S2A**).

### NMF captures common metagenes that strongly contribute to expression patterns of immune checkpoint modulators

To explore underlying patterns between the response groups while reducing statistical overfitting, we applied a nonnegative matrix factorization (NMF) technique (Lee and Seung, 1999) for clustering and metagene extraction in the merged dataset and two individual datasets. At this point, we used subsets of the gene expression matrices of the common genes selected from LASSO. Hierarchical clustering showed no evidence of a potential association between NMF results and response or putative confounders (*p* = 0.2549, Fisher’s exact test; **Figure 2A**). Of note, the three sets of NMF results manifested completely concordant “metagene determinants” (*CEACAM1, CD40, B7-H3*, and *CD112*) at rank = 4; the optimal rank was selected according to the NMF rank survey (**Figure S3A**). Such metagene determinants indicate that the genes with the strongest contribution to each cluster (i.e., metagene) are consistent across the merged dataset (**Figure 2B**) and the two individual datasets (**Figure S3B**) (empirical *p* = 0.001). Thus, this concordance implies that the metagene determinants may play important roles in melanoma immune status, and the two melanoma datasets share an analogous orchestration of the expression of immune checkpoint modulators.

**Figure 2.**
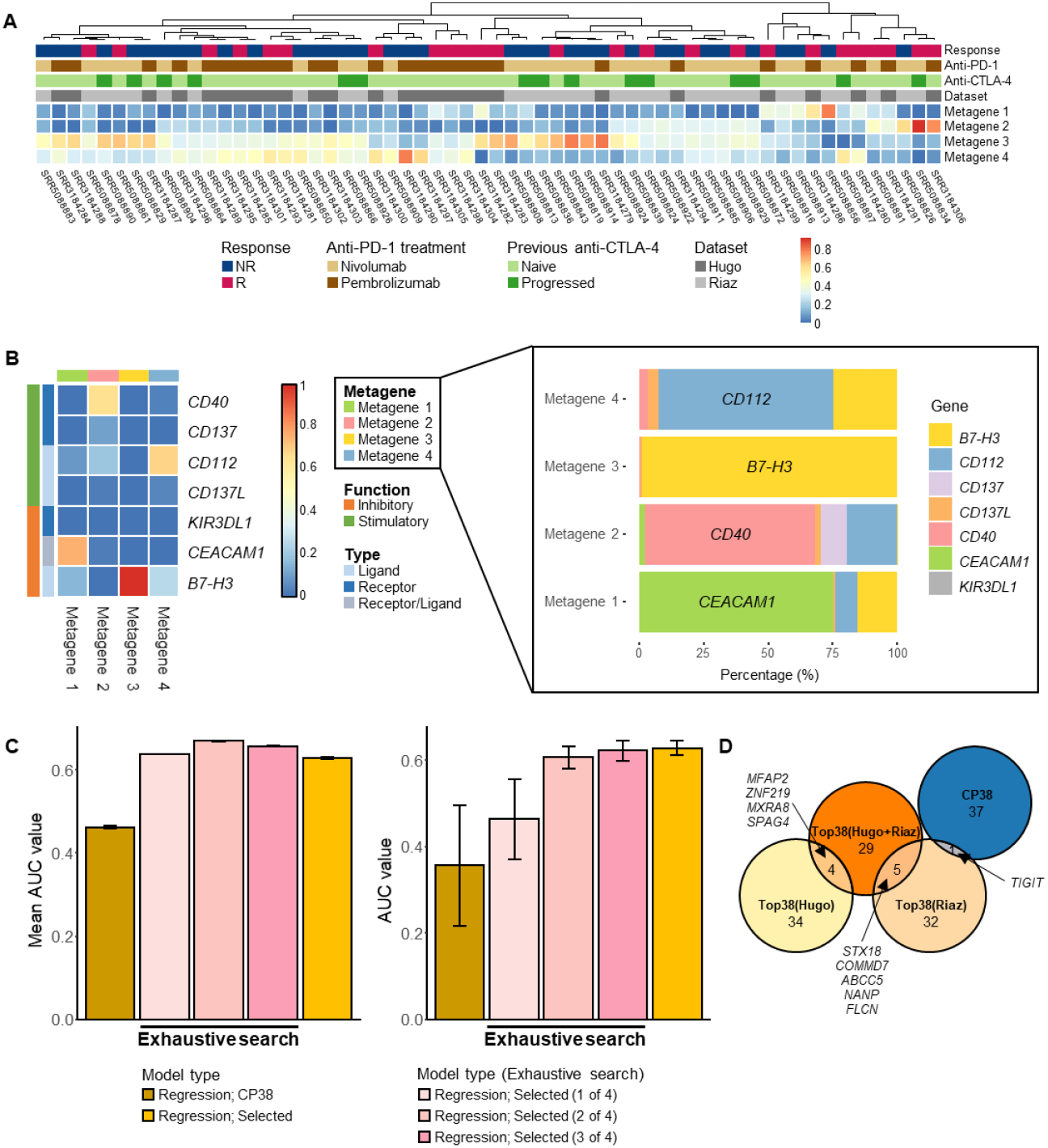
Results from the regression-based approach. (A) Matrix H obtained from NMF. This heatmap depicts the contributions of each metagene to the individual melanoma patients (n = 60). Each column is scaled to sum up to one; the red color indicates a higher contribution of each metagene. (B) Matrix W obtained from NMF. This heatmap (**left**) depicts the contributions of each gene to 4 metagenes. The stacked bar plot (**right**) shows the graphical illustration of the composition of each metagene. The metagene determinant that contributed most strongly to each metagene is as follows: Metagene 1 (*CEACAM1*), Metagene 2 (*CD40*), Metagene 3 (*B7-H3*), and Metagene 4 (*CD112*). (C) Mean AUCs of both 5- and 10-fold CV: exhaustive regression models (without including the *TDS*) (**left**); AUCs from vice versa prediction using exhaustive search-based regression (without including the *TDS*) (**right**). (D) Euler diagram of the 38 top-ranked DEGs (i.e., Top38) selected in different datasets, along with the 38 immune checkpoint modulators (i.e., CP38) selected in the merged dataset. The top-ranked DEGs were selected in each dataset: H indicates Hugo et al. (Hugo et al., 2016) (light yellow), R indicates Riaz et al. (Riaz et al., 2017) (light orange), and HR indicates the merged dataset (orange). CP38 indicates the 38 immune checkpoint modulators selected in the merged dataset (blue).

### Exhaustive search-based regression selects the best subsets among the metagene determinants

The predictive power of 10-fold cross-validation (CV) using a regression model with the TPM values of the basal gene set was essentially nonsignificant (mean area under the curve (AUC) = 0.46). Conversely, the common metagene determinants from NMF showed improved predictive power (mean AUC = 0.63). We further demonstrated an exhaustive search using the 4 metagene determinants selected via the regression approach based on the Akaike information criterion (AIC) and Bayesian information criterion (BIC) (**Figures S2B-C**). As a result, we selected the best subset of *CEACAM1* and *B7-H3* that showed the highest performance at a mean AUC of 0.67 from both 5- and 10-fold CV (**Figure 2C (left)**). The vice versa prediction (see **Methods**) showed lower performance (mean AUC = 0.57), which indicates putative heterogeneity between the two datasets (**Figure 2C (right)**). Additional investigation of differentially expressed genes (DEGs) in each dataset and the merged dataset supported such heterogeneity. When the same number of DEGs (i.e., 38) was selected as the basal gene set based on the lowest *p* values from each dataset, no genes overlapped between the two individual datasets, and only 4 and 5 DEGs from the combined dataset overlapped with those from each dataset (**Figure 2D**).

### Correlations of immune checkpoint modulator expression manifest striking differences according to anti-PD-1 responses

First, we observed that the correlations among the TPM values of the 38 immune checkpoint modulators showed discriminative patterns for anti-PD-1 responsive melanoma (**Figure 3A**). These differential patterns suggest that the expression levels of these genes vary according to the response. The responders showed strong positive correlations between the expression levels of the immune checkpoint modulators, implying that they are activated in the immune system, whereas the nonresponders showed weaker correlations. Accordingly, the overall distribution of correlations between responders and nonresponders showed a substantial difference (*p* = 4.44×10^−15^, Kolmogorov-Smirnov test; **Figure S4**). Altogether, these correlation patterns are consistent with the concept of hot and cold tumors (Galon and Bruni, 2019), which determines responsiveness to ICIs in cancer patients.

**Figure 3.**
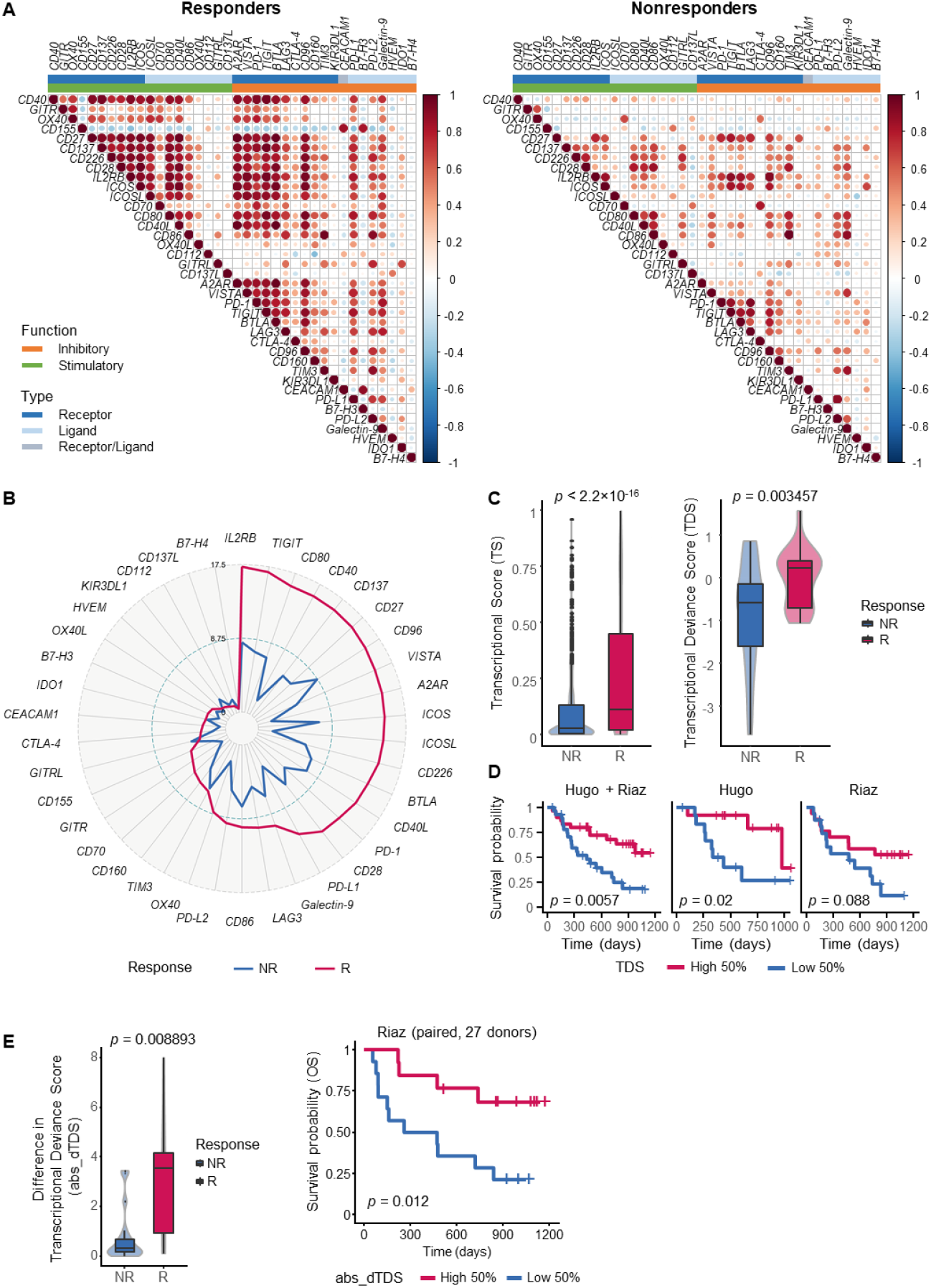
Results from the correlation-based approach. (A) Correlograms showing correlations between the 38 immune checkpoint modulatory genes in each response group (responders (**left**) and nonresponders (**right**)) to PD-1 blockade. (B) The radar plot of the sum of the coefficient of determination (*R*^*2*^) by the response group. The sum of the *R*^*2*^ values comprises the *TS*. Genes are ordered by the *R*^*2*^ values of the responders, in descending order. (C) Box plots presenting each *R*^*2*^ value (**left**) and *TDS* (**right**); the *TDS* represents the individual-wise contribution to the *TS*. The violin plot that is merged with each box plot depicts the distribution of the values. (D) Kaplan-Meier plots of OS for three dataset configurations (Hugo + Riaz, Hugo, and Riaz). All plots were divided by the *TDS* status (high or low) by binarizing the *TDS* with the dataset-wise median, and *p* values of the likelihood ratio test (LRT) in the Cox Proportional-Hazards model fit are presented. (E) Box plots showing the absolute values of the difference in the *TDS* by the response (**left**) and Kaplan-Meier curves (**right**) for 27 paired donors included in the Riaz et al. dataset (pre- and on-therapy samples); *p* values from two-sided Wilcoxon test and log-rank test are presented, respectively.

### Introducing the *TS* to summarize immune engagement based on the expression of immune checkpoint modulatory genes

Herein, we introduce an intuitive correlation-based measure, the *transcriptional score (TS)*, to capture both an overall difference between the two response groups and relevant synergistic effects within the basal gene set. In short, the *TS* is a gene-wise measure that sums up all the coefficients of determination (*R*^2^) between a gene and the rest of the genes within our basal gene set. In this respect, the *TS* reflects the degree of explanatory power between each gene and the remaining genes (see **Methods**). Despite its conciseness, our results show that the *TS* vividly distinguished the responders from the nonresponders (**Figure 3B**). Moreover, 66% (25 out of 38) and 92% (35 out of 38) of the genes in the basal gene set showed statistically significant segregation according to the *TS* and an elevated *TS* in the response group, respectively (**Figure S5**). Specifically, *IL2RB* showed the highest *TS* (17.24), while *B7-H4* showed the lowest (0.44) (**Figure 3C (left)**). Furthermore, the individual *R*^*2*^ values that comprise the gene-wise *TS* showed evident segregation by the response group (*p* < 2.2×10^−16^, Welch’s *t*-test; **Figure S6A**).

Additional simulation (see **Methods**) revealed that the *TS* ratio, which is a ratio of responders’ *TS* to nonresponders’ *TS*, is likely to originate from the normal distribution, which is highly beneficial for identifying its statistical properties. Moreover, many of the gene-wise *TSs* were statistically significant (minimum empirical *p* <0.0006, 7 genes; **Figure S6B**) and enriched (minimum empirical *p* < 4.13×10^−5^, 14 genes; **Figure S6C**), and construction of the background distribution revealed that the basal gene set had profound background differentiation and relative enrichment (*p* =2.37×10^−7^ and 1.98×10^−8^, Wilcoxon’s rank-sum test; **Figures S6B-C**).

### The *TDS* serves as a powerful sample-wise predictor derived from the *TS*

Additionally, we defined quantifiable individual-wise contributions in the construction of the *TS* as a new concept named the *transcriptional deviance score (TDS)*. In short, the *TDS* for the particular *i*^th^ individual is calculated by subtracting the *TS* that corresponds to the rest of the samples without the *i*^th^ sample from that corresponds to the entire sample set (see the illustration in **Figure S2D**). Hence, this calculation measures the individual contribution of the *i*^th^ sample to the overall *TS* of the given sample set (see **Figure 1C** and **Methods**). Similar to the *TS* values, the *TDS* values were significantly elevated in the responders (*p* = 0.003457, Welch’s *t*-test). This implies that the responders are likely to show the orchestration of immune checkpoint modulators (**Figure 3C (right)**).

Using the *TDS*, we replicated the exhaustive search performed in the regression modeling (**Figure S2E**). Consequently, the model including three variables (the TPM values of *CEACAM1* and *B7-H3* and the *TDS*) showed the highest mean AUC of 0.74, which was improved from the mean AUC of 0.67 by adding the *TDS* to the model. Hence, this reveals the *TDS* as a potentially effective variable to strengthen the model performance. Accordingly, the three-variable model including the TPM values of *CD40* and *CD112* and the *TDS* presented the highest AUC of 0.81 when trained on the Riaz et al. (Riaz et al., 2017) dataset and tested on the Hugo et al. (Hugo et al., 2016) dataset. In addition, implementing the exhaustive searches under the two conditions (i.e., with the *TDS* and without the *TDS*) enabled us to confirm the advantage of including the *TDS* in the final subset.

Moreover, regardless of the dataset, survival analyses using the *TDS* revealed its powerful ability to segregate survival status (**Figure 3D**). In both the combined and individual datasets (see **Methods**), overall survival (OS) was separated by the median *TDS*, which was calculated without any phenotypic information but with only the gene expression of the immune checkpoint modulators (*p* = 0.0057, 0.02, and 0.088 for the combined, Hugo et al., and Riaz et al. (Riaz et al., 2017) datasets, respectively, log-rank test; **Figure 3D**), and the *p* value was further improved in the combined dataset. However, the DEGs related to the OS status from the individual datasets did not overlap, and only a small fraction of genes were commonly identified between the combined and individual datasets (*TPD52* and *MYO5B* from the Hugo et al. dataset and *EPS8L1* and *NAPB* from the Riaz et al. dataset).

### Dynamic changes in the *TDS* following ICI treatment correlate with ICI responses

We evaluated whether the *TDS* reflects how the dynamics of immune networks are affected by ICI therapies. In this respect, we compared the changes in the *TDS* after ICI treatment using pre- and on-treatment samples collected from 27 donors who were included in the Riaz et al. (Riaz et al., 2017) dataset. We calculated the *absolute values of differences in the TDS (abs_dTDS)*. As a result, we confirmed that the *TDS* values of the pre- and on-therapy paired samples show significant differences (*p* = 0.008893, two-sided Wilcoxon test; **Figure 3E (left)**) according to the response to ICIs. Furthermore, the differences in the *TDS* following ICI treatment significantly segregated the OS status (*p* = 0.012, log-rank test; **Figure 3E (right)**).

### The predictive power of the *TDS* is strengthened by a combination of existing predictive markers

We established a multitude of predictive models using various sets of genes that were identified via both regression- and correlation-based approaches. To compare the predictive power, we included 4 FDA-approved immuno-modulatory target genes (*PD-1, PD-L1, PD-L2*, and *CTLA-4*) as reference genes in the evaluation. For a concrete comparison, we generated 4 models covering either single (*PD-1* or *PD-L1*) or multiple (*PD-1*+*PD-L1*+*CTLA-4* or *PD-1*+*PD-L1*+*PD-L2*+*CTLA-4*) targets as predictors.

Throughout predictive modeling with the two approaches, the predictive performance was substantially improved compared to that of the model using the conventional biomarkers (**Figure 4A**). Consistently, the CV (**Figure 4A (top)**) and train-test (**Figure 4A (bottom)**) schema exhibited similar patterns. Specifically, the predictive performance was higher in both the regression-based model (AUC = 0.63) and the correlation-based model (AUC = 0.71), whereas the models designed based on the conventional biomarkers showed the lowest value (AUC = 0.33). Notably, the combined model of the two genes (*CEACAM1* and *B7-H3*) from the regression-based approach and the *TDS* from the correlation-based approach enhanced the performance, with the highest AUC of 0.75 (**Figure 4A (top)**); when the four metagene determinants were added to the models, the performances were moderately high (AUC above 0.7) but did not reach the highest AUC of the model from the exhaustive search. Additionally, our proposed models showed smaller differences in performance between the training and testing datasets (**Figure S7A (left)**) or the 10-fold (**Figure 4A (top)**) and 5-fold CV sets (**Figure S7A (right)**). This confirms that our approaches to data processing and modeling successfully reduced any potential prediction biases. Moreover, the vice versa prediction (see **Methods**) verified that our proposed models demonstrated a subtle range of fluctuations in AUCs, indicating that the performance of the model was not hindered by the heterogeneity of the datasets used (**Figure 4A (bottom)**). Meanwhile, we did not consider race-based disparities in the therapeutic responses to PD-1 inhibition, due to there being insufficient data for each donor included in the dataset used in this study.

**Figure 4.**
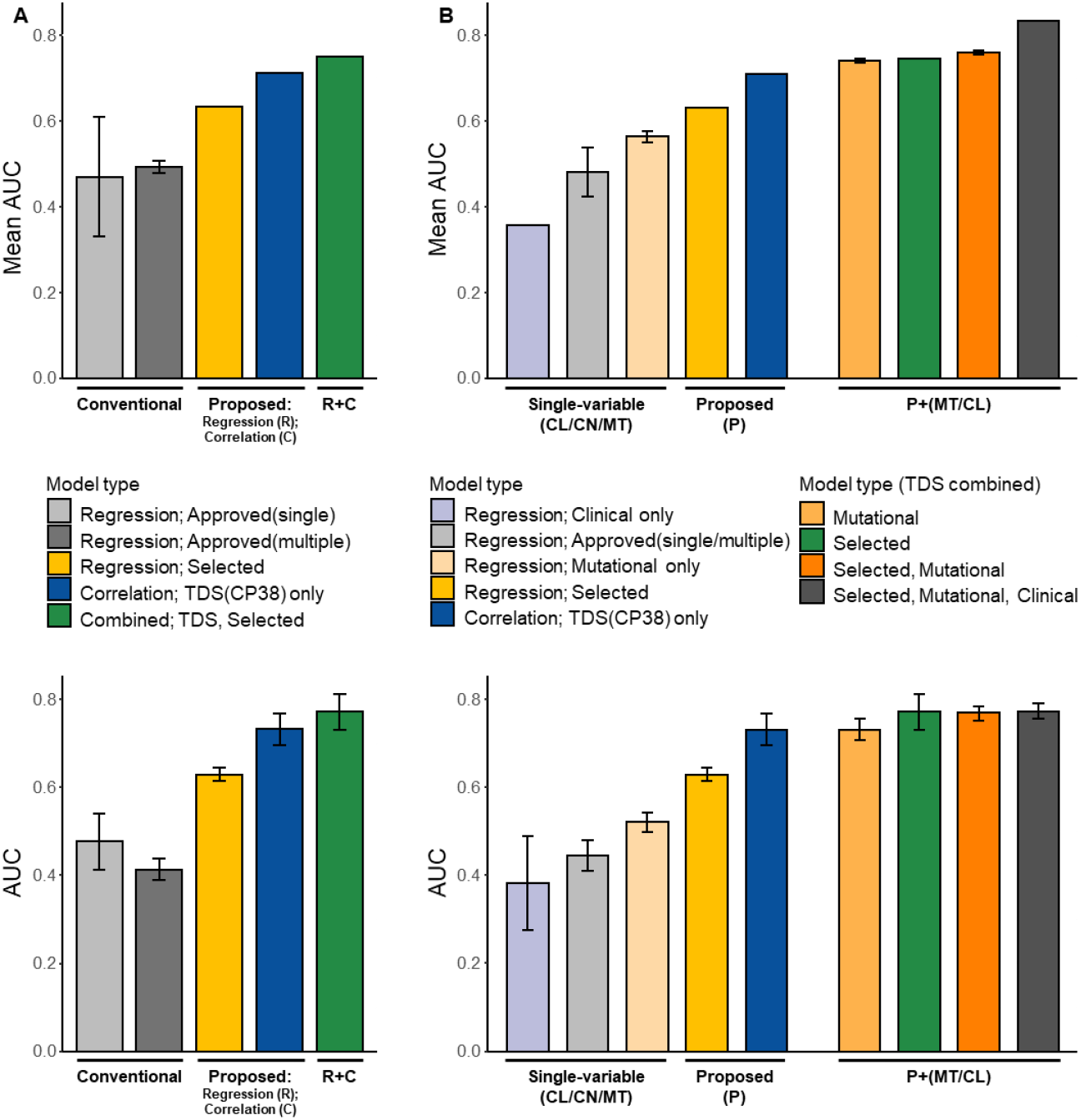
Model performance evaluation. (A) Bar plots of the mean AUCs from 10-fold CV (**top**) and the AUCs from the vice versa train-test scheme (**bottom**). These bar plots cover the models established with approved targets (i.e., Conventional), regression (R) or correlation (C) approaches (i.e., Proposed), and combined approaches (i.e., R+C). (B) An extensive summary of the performance evaluation of different models showing the mean AUCs from 10-fold CV (**top**) and the AUCs from the vice versa prediction (**bottom**). These bar plots cover the models established with single variables (either clinical (CL), conventional (CN), or mutational (MT) variables), the proposed variables, and the combined variables. The clinical variable refers to the metastasis stage (M stage), and the mutational variable refers to triple wild-type status for significant mutations (i.e., *NRAS, NF1*, and *BRAF*) in melanoma.

To strengthen the model performance, we added the binary mutation status (*NRAS, NF1, BRAF* mutant, or triple wild-type status) and polynomial metastasis stage (M stage) classification of the individual patients to our proposed models. The combined models thereby outperformed the 10-fold CV (repeated 100 times), with an AUC of 0.83 when the two variables (triple wild-type and M stage) were added to the exhaustive search-based model including the *TDS* (**Figure 4B (top)**), and correspondingly showed high performances with the vice versa prediction (**Figure 4B (bottom)**). In summary, the final model includes 5 variables: (1) *TDS*; (2) M stage category; (3) binary status of triple wild-type or *NRAS, NF1*, and *BRAF* mutations; (4) gene expression levels of *CEACAM1*; and (5) gene expression levels of *B7-H3*.

### Comparison of model performance confirms the superiority of *TDS*-based prediction models

For further model evaluation, we compared predictive performance between the model developed with the *TDS* calculated from the 38 top-ranked DEGs and our model (comprising the basal gene set). Consequently, the *TDS*-based model constructed with the basal gene set showed better performance (mean AUC = 0.73) than that constructed with the top DEGs (mean AUC = 0.56) in the vice versa train-test evaluation scheme (**Figure S7B**). Thus, the differences between the AUCs indicate that the predictive power of the *TDS* does not rely upon the selection of marginally significant genes in ascending order of *p* values. Additionally, vice versa predictions using the top-ranked DEGs selected from the combined dataset revealed the predictive performance of each DEG. The individual predictions were performed in the combined dataset of Hugo et al. and Riaz et al. and the individual datasets and thereby demonstrated unstable and poor performance (**Figure S7C**). Next, we consolidated the power of including *TDS* in the models by comparing the models developed with both regression and correlation frameworks (**Figure S7D**). First, we compared the models established based on expression values (in TPM) of our basal gene set with the model established based on the expression values of the 4 selected immune checkpoint modulators via regression-based feature selection. This comparison indicated that our regression approach had favorably chosen genes that may be associated with determining the response. In addition, the performance of the models developed with the expression of the 4 selected genes surpassed that of the models developed with the TPM values of the 38 top-ranked DEGs and showed power equivalent to that of the models based on the *TDS* calculated from the same sets of the top-ranked DEGs. Last, we compared the correlation-based model using the basal gene set with the artificially designed models using 38 randomly selected genes (with 10 repetitions). Hereby, we noted that the correlation-based model (with *TDS* of the basal gene set) outperformed (mean AUC = 0.71) the random selection-based artificially designed models (mean AUC = 0.5). This confirms the importance of including biologically meaningful genes involved in the immune checkpoint modulatory system in the model, which is conveyed by the impact of the *TDS*. The regression-based model with the metagene determinants, however, demonstrated improved predictive performance compared to that of the model based on the expression of the basal gene set. Moreover, the metagene determinant-based model showed predictive power equivalent to that of the model including the 38 DEGs, demonstrating the validity and parsimony of our regression-based variable selection method.

Model development based on the M stage magnifies the impact of including the selected variables from our proposed approaches, as the mean AUCs were augmented after these variables were added into the single-variable model including the M stage only (**Figure S7E**). This corroborated the impact of using the *TDS*, which conveys both statistical and biological insights, in the predictive model. In summary, performance evaluation proved the strong contribution of the *TDS*, which includes the immune checkpoint modulators that were showed varying basal expression levels (**Figure S8**).

### Integration of the biological domain knowledge-driven regression and correlation analytic frameworks selects balanced features representing the immune status

With the integration of regression-based and correlation-based approaches, we summarized the key findings into the transcriptional determinants for PD-1 inhibition (**Figure 5A**). We further evaluated the ratio of the *R*^*2*^ values in responders and nonresponders (**Figure 5B**) and identified the genes that showed distinctive features. Notably, we found that the 4 genes selected via the regression-based approach were distributed evenly among high (8.52, *CD40*), medium (3.90, *CEACAM1*), and low (2.39, *B7-H3* and 0.83, *CD112*) ratios; these genes act as a stimulatory receptor, an inhibitory receptor/ligand, an inhibitory ligand, and a stimulatory ligand, respectively. We thus infer that our regression method selected the optimal genes that represent each role and function and are response-dependent. We used a differentiated strategy of regression-based variable selection to develop the model with the selected genes, which exhibited predictive power equivalent to that of the model developed with the DEGs and performance superior to that of the model developed using our basal gene set (**Figure S7D**).

**Figure 5.**
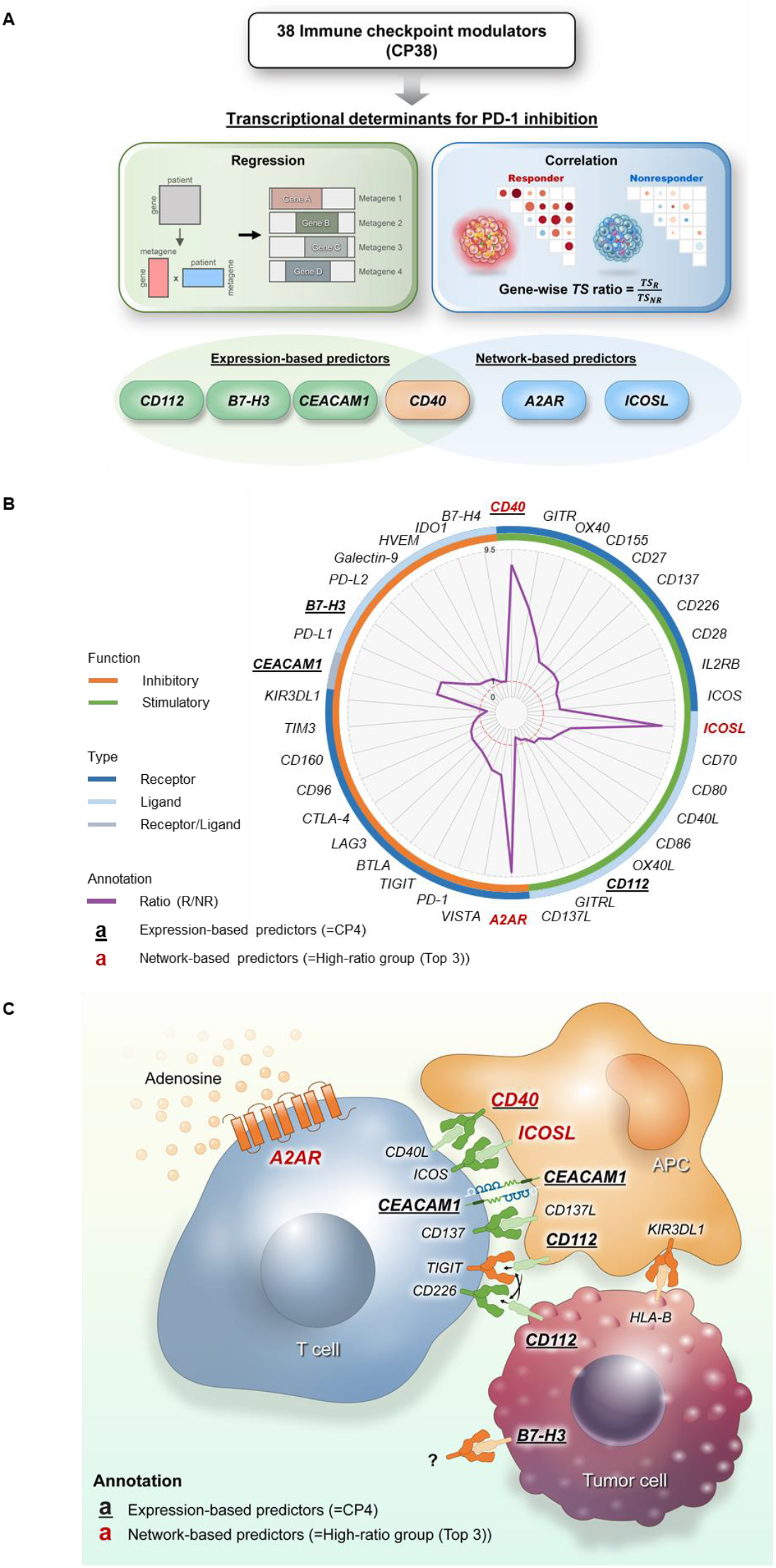
Significance of our proposed models. (A) Graphical diagram summarizing the major findings from the two approaches: the transcriptional determinants for PD-1 inhibition selected via the regression- and correlation-based frameworks. (**Left**) Based on the regression approach, expression levels of the 4 genes (*CD112, B7-H3, CEACAM1*, and *CD40*) were selected. (**Right**) Based on the correlation approach, networks driven by the three genes (*CD40, A2AR*, and *ICOSL*) were selected. The simplified illustrations highlight the general idea of how these two approaches were performed to select the transcriptional determinants. (B) Radar plot of the ratio of the sum of the *R*^*2*^ values calculated for each of the 38 immune checkpoint modulators in the responders to nonresponders. The genes selected via the regression approach are underlined. Genes are ordered by the ratio in descending order and partitioned according to immunological categories based on both function and type: functions include stimulatory and inhibitory and types include receptor, ligand, and receptor/ligand. The top-ranked genes (written in red) were determined as *A2AR* (8.97), *ICOSL* (8.46), and *CD40* (8.20); the numbers written in the parentheses indicate the ratio of the values in the responders to nonresponders. (C) Biological expression sites of the selected immune checkpoint modulators among the basal gene set. Binding between each ligand and receptor is illustrated.

The *TDS* conveys the association between the 38 immune checkpoint modulators and denotes prominent predictors to distinguish underlying immunological parameters in melanoma patients. Based on the *TDS*, our final model outperformed by including combined features (such as gene expression levels, mutational status, metastasis classification, and correlation-based interaction scores) that depict the immunotherapeutic potential of melanoma tumors. The genes located in the high-ratio group (i.e., *A2AR* (9.18), *ICOSL* (8.66), and *CD40* (8.52)) likely show significant differences in gene expression levels between responders and nonresponders (**Figure 5B**).

Based on the ratio rank, *A2AR* showed the highest ratio, with a 9-fold higher sum of *R*^*2*^ values in responders than in nonresponders. In other words, *A2AR*-centered interactions with each of the remaining of the 37 immune checkpoint modulators were more strongly activated in anti-PD-1 responders than in nonresponders; in particular, the interactions between *A2AR* and *ICOSL* (the second-highest gene) were statistically significant (*p* = 0.0463, binomial generalized linear model). Furthermore, the immune checkpoint modulators selected with the two approaches were depicted by the expression sites: a tumor cell, T cell, or antigen-presenting cell (APC) (**Figure 5C**).

### Estimation of immune cell fractions does not distinguish anti-PD-1 responders from nonresponders

To characterize immune cell types, we estimated the fractions of 22 types of immune cells (LM22) by employing CIBERSORTx (Newman et al., 2015; Newman et al., 2019) (**Figure S9**). We thereby noticed that the fractions of M2 macrophages were very large in the overall population of melanoma patients (**Figure S9A**). In addition, only three types of immune cells showed subtle differences in the ratio of the sum of *R*^*2*^ values according to anti-PD-1 response (above a ratio of 2): memory B cells (3.68), gamma delta T cells (2.65), and neutrophils (2.26). Nevertheless, unlike the distribution of the *TDS*, the fractions of the immune cells did not show distinct differences between the response groups (**Figure S9B**). The fractions of these three immune cell types showed poor predictive performance (range of AUCs = 0.47-0.57 (10-fold) and 0.45-0.54 (5-fold)).

### Homologous protein sequences of B7-H3 and PD-L1 suggest shared functional roles in predicting anti-PD-1 responses

The 4 selected genes from the regression method included neither *PD-1* nor *PD-L1*, which serve as the major approved targets for immunotherapy. Thus, to discover specific genes that may play alternative biological roles in the PD-1/PD-L1 axis, we conducted sequence alignment analyses by using the protein-protein Basic Local Alignment Search Tool (BLASTp) to investigate the sequence homology of PD-1 or PD-L1 with each selected gene. From the protein-level sequence alignment, we observed that the sequence alignment between B7-H3 and PD-L1 yielded the highest total alignment score (257) among the alignment combinations. Specifically, the highest total score mainly encompasses the maximum scores yielded from the alignment of two sequence regions (17 to 288 (max score = 126) and 12 to 288 (max score = 113)) with sequences that belong to the extracellular domain of PD-L1 (**Figure S10** and **Table S3**).

## Discussion

Two approaches were implemented in this study based on regression and correlation frameworks to improve our biological understanding of anti-PD-1 responses using transcriptomic datasets from melanoma patients. Using our basal gene set, leading genes and indices of immune status were simultaneously selected via regression-based (**Figure 1B**) and correlation-based (**Figure 1C**) approaches, respectively. Both approaches included a novel concept to avoid overfitting issues from considering the phenotypes (i.e., responses to PD-1 inhibitors). The regression approaches elicited the 4 key metagene signature determinants through NMF, which utilizes the phenotype-independent method; this approach also enabled us to obtain biological insights. Through the correlation approach, the final predictor, the *TDS* (**Figure 3C (right)**), was established solely based on the expression values of the immune checkpoint modulators. Despite the exclusion of phenotypes, our biological knowledge-driven approaches successfully achieved predictive power superior to that of any other prediction models with known biomarkers. Additionally, no significant race-based disparities were found in anti-PD-1 responses according to a previous study (O’Connor et al., 2018). Thus, our study confirmed the association between the immune status and ICI responses, which are obvious biological consequences, and indicated that these approaches can be used to elucidate the immune status in melanoma.

First, the predictive model established based on the expression of the 4 genes selected by the regression approach improved predictive performance compared to that of the two models developed based on the statistically significant gene sets and the basal gene sets. Even the combination of the selected predictors progressively augmented predictive power for differentiating responses in melanoma (**Figure 4**). Conversely, this consistent and outstanding predictive performance was not observed when the models based on either the top-ranked DEGs (**Figure S7C**) or randomly selected genes were used. These models showed variable performance. In particular, the model including only the *TDS* of the basal gene set (*TDS*-only model) stably outperformed the regression-based and artificially designed *TDS*-based models (**Figure S7D**). These results confirm that the genes selected via the regression approach act as significant predictors of immune status in melanoma.

Moreover, *TS* is the sum of coefficients of determination (*R*^*2*^) and represents the gene-gene interactions among the 38 immune checkpoint modulatory genes, based on magnitude only, without regard to the direction of each correlation. This score was designed to quantify the overall degree of the immune network or engagement in the tumor microenvironment of anti-PD-1-treated melanoma patients. Most importantly, the *TDS*, a novel score reflecting individual-wise immune engagement, showed excellent correlations with therapeutic responses (**Figures 3C (right)**) and predicted response when included in the models (**Figure 4B** and **S7D**). Specifically, the predictive power increased progressively after the *TDS* was added to the other models that used single variables such as mutational and clinical features (**Figures 4A-B**). When these features were included alone, the performance was poor (AUCs below 0.6). Unlike the *TDS*, the immune cell fractions of individual patients did not show distinct patterns according to the response (**Figure S9B**) and showed unstable, poor performance in predictive modeling. In addition, the DEG-based models demonstrated fluctuations in performance rates, and the DEGs themselves did not provide interpretable results as they rely on statistical significance only, not biological significance (**Figures S7B-C**). We repeatedly confirmed the superiority of the *TDS*, as it reflects the temperature of immunological engagement (hot and cold tumors (Galon and Bruni, 2019)), which many researchers are eager to delineate; hence, we suggest this score as a powerful indicator for understanding the tumor microenvironment. Furthermore, we compared the changes in the *TDS* of the paired sample sets — pre- and on-therapy samples — of 27 donors included in the Riaz et al. (Riaz et al., 2017) dataset (**Figure 3E**). This result demonstrates that the dynamics of the correlation-based immunobiological networks are more activated in individuals who respond to PD-1 inhibition in melanoma.

In summary, our combined approaches using the regression and correlation frameworks successfully achieved stable predictive power for distinguishing responses in melanoma. For further application, this new concept can be utilized for both developing prediction models and deducing biological interpretations in any cohort, regardless of the population size.

TMB has been considered an independent predictive biomarker in various cancers treated with immunotherapy (Goodman et al., 2017; Rizvi et al., 2015; Rosenberg et al., 2016; Snyder et al., 2014; Van Allen et al., 2015). Unlike other cancer types, melanoma shows distinct genomic features across subtypes (Hayward et al., 2017). Accordingly, TMB alone does not correlate with ICI responses (Keenan et al., 2019), and the association between TMB and response is confounded by subtype (Liu et al., 2019). In summary, TMB cannot predict response in melanoma, which is supported by a recent review (Luke and Ascierto, 2020) stating that the median TMB was higher in responders and simultaneously overlapped with nonresponders. Hence, considering the incompatibility of TMB with melanoma subtype, we note that our proposed models including the *TDS* outperformed the other models in predicting response, without involving TMB. Furthermore, we suggest that a combination of the *TDS* with TMB might be effective in predicting response in other cancer types.

The final model includes 5 predictors: (1) *TDS*, (2) categorical M stage, (2) the binary status of the wild-type status of the three mutations (in *BRAF, NRAS*, and *NF1*), and expression of (4) *CEACAM1* and (5) *B7-H3*. Collectively, each predictor contributed connections with both phenotypes and genotypes of melanoma patients. Specifically, *CEACAM1* is involved in intercellular cell adhesion, which varies by expression site. According to recent studies, the high expression of *CEACAM1* promotes the progression and metastasis of melanoma (Turcu et al., 2016; Wicklein et al., 2018). *B7-H3* (*CD276*) contributes to the evasion of immune surveillance and cancer progression (Cai et al., 2019; Castellanos et al., 2017; Yonesaka et al., 2018). Furthermore, dual blockade of B7-H3 and PD-1 effectively reduced tumor size, indicating that B7-H3 and PD-1 are promising candidates for immunotherapy (Lee et al., 2017). In summary, these functional roles of both *CEACAM1* (Dankner et al., 2017) and *B7-H3* (Picarda et al., 2016; Robert et al., 2011) may be associated with the prediction of anti-PD-1 responses in melanoma.

According to a further comparison with other approaches that utilize the same datasets, Auslander et al. suggested that the immuno-predictive score (IMPRES) is a new transcriptional predictor of responses to ICIs (Auslander et al., 2018) by demonstrating that its prediction performance rates are the most powerful of the existing methods. Until recently, the validity of the concept used in the IMPRES was still actively debated (Auslander et al., 2019; Carter et al., 2019), and hence, we anticipate an active comparison with the concept used in our model that was developed based on the *TDS*. As a result of the comparison, our approach achieved a superior predictive power compared to that of the existing method (AUC = 0.62 in HR60, 0.7 in Hugo27, and 0.58 in Riaz33 datasets).

Additionally, the three genes that showed the highest ratios of the sum of *R*^*2*^ values in responders (*A2AR, ICOSL*, and *CD40*) were associated with ICI responses. First, *A2AR* (*ADORA2A*, an adenosine A2A receptor) binds to extracellular adenosine (ADO), a metabolite of ATP, which accumulates in the hypoxic tumor microenvironment (Ohta, 2016; Vigano et al., 2019). Recent studies described the pleiotropic effects of adenosine and different outcomes of A2AR deficiency in the tumor microenvironment according to tumor type (Cekic and Linden, 2014; Vigano et al., 2019). In addition, A2AR signaling triggers the expression of PD-1 on effector T cells, which supports the synergistic effect of recent combination therapy trials of A2AR antagonists with PD-1 inhibitors in cancer treatment (Beavis et al., 2015; Leone et al., 2018; Mittal et al., 2014; Sek et al., 2018). Second, *ICOSL* (a ligand of *ICOS*) plays dual antitumor and protumor roles by regulating tumor activities when bound to ICOS (Solinas et al., 2020). Solinas et al. (Solinas et al., 2020) emphasize that further studies are required to understand how ICOS-ICOSL interactions contribute to triggering memory immune responses of CD4+, CD8+, and B cells in tertiary lymphoid structures (TLSs). In addition, the activation and expansion of regulatory T cells were enhanced in melanoma cells expressing ICOSL (Martin-Orozco et al., 2010). In particular, the functional roles of ICOS-ICOSL substantiate combination therapies with either PD-1 or CTLA-4 inhibitors in recently ongoing clinical trials (Solinas et al., 2020). Last, *CD40* is involved in antigen presentation and B cell activation under T cell-mediated conditions. Recent studies demonstrated the effective antitumor immunity induced by CD40-activated B cells, which trigger T cells to respond, and highlighted the potential of B cells in cancer immunotherapy (Guo and Cui, 2019; Kornbluth et al., 2012; Wennhold et al., 2019). Additionally, the presence of B cells in TLSs was frequently observed in tumor cells with higher response rates to immunotherapy. Furthermore, increased expression of *CD40* indicates more mature TLS signatures (Cabrita et al., 2020; De Silva and Klein, 2015), leading to a correlation with survival benefits in cancer patients (Sautes-Fridman et al., 2019). These findings confirm the pivotal roles of B cells present in tumor cells in inducing optimal antitumor responses to immunotherapy.

Furthermore, we found that *B7-H3* may have potentially redundant biological functions with *PD-L1* (*CD274* or *B7-H1*), as these two genes both belong to the B7 superfamily, which is involved in regulating T cell responses (Ni and Dong, 2017). According to previous studies (Chapoval et al., 2001; Ni and Dong, 2017), B7-H3 has sequence identity with the extracellular domain region of PD-L1, which corresponds to our BLASTp results. These findings confirm that these two genes may share inhibitory functions based on the homologous region, where binding of receptors and ligands and docking of inhibitor molecules occur. Collectively, the previous functional studies showed that both genes are involved in inhibiting T cell proliferation (Dong et al., 2002; Suh et al., 2003; Veenstra et al., 2015), leading to aggressiveness and proliferation in tumor cells and thereby mediating immune evasion (Castellanos et al., 2017; Flem-Karlsen et al., 2019; Picarda et al., 2016; Tekle et al., 2012).

It is notable that *CD40* was selected by both approaches. This highlights the importance of B cell-mediated immunotherapy in triggering T cell responses, underlying the potential biological roles of *CD40*. Thus, we suggest that CD40-activated immune status might reflect the network effect of immune checkpoint modulators and represent the responsiveness to ICIs in melanoma.

In conclusion, our findings summarize the RNA-based ICI-related biomarkers that simultaneously predict both responses and survival. Based on the hot and cold tumor concept, we validated that the *TDS* indeed represents immune phenotypes, especially immune temperatures, which can be interpreted by the summation of the networks established in the tumor and immune cells. Additionally, we examined the *TDS* distribution in lung squamous cell carcinoma (LUSC), which has a very similar carcinogenic mechanism to melanoma (carcinogen-driven cancer) and rather homogenous phenotypes among lung cancers. Interestingly, LUSC patients manifested a comparable *TDS* distribution pattern, as the *TDS* seems to be higher in ICI responders than nonresponders (p = 0.1286, one-sided Wilcoxon rank-sum exact test; **Figure S11**). This approach suggests that the *TDS* can be applied in other cancer types. Hence, further investigations will enable us to pinpoint potential networks among immune checkpoint modulators or biologically related genes in other cancer or disease datasets with the adequate adjustment and refinement of primary gene sets for the relevant cancer subtype and therapeutic axes. In addition, our study will be strengthened if new immune checkpoint modulatory genes are discovered and the networks among them are uncovered. Further plans for *in vitro* experiments using retrospective clinical specimens from melanoma patients and validation in larger cohorts of patients with different cancers are being considered to strengthen these findings with comparative studies; scarce resources with transcriptomic data paired with anti-PD-1 responses and clinical parameters were available publicly. In summary, our integrative analytical workflow, which is a combination of traditional and novel approaches based on domain knowledge, is flexible and can be applied to investigate the immune status of cancer patients and to discover surrogate biomarkers for predicting therapeutic responses and broadening clinical benefits.

## Acknowledgments

This study was supported by the Basic Research Laboratory (BRL) program through the National Research Foundation (NRF) funded by the Ministry of Science and ICT (MSIT) (grant accession: 2018R1A4A1022513), and supported by the Korea Health Technology R&D Project through the Korea Health Industry Development Institute (KHIDI), funded by the Ministry of Health & Welfare (grant accession: HI14C1277). We thank the Global Science Experimental Data Hub Center (GSDC) and the Korea Research Environment Open NETwork (KREONET) service for data computing and the Korea Institute of Science and Technology Information (KISTI) for providing these networks.

## Author Contributions

H.S. and Y.K. conceived and designed the overall study. H.S. contributed to data collection, analysis, and visualization. S.L. contributed advice and assistance related to the statistical and analytical aspects of this study. H.S. and S.L. contributed to manuscript writing via discussions with Y.K. and H.Y. The validation data generated from clinical specimens were provided by S.-H. L. M.K. provided clinical perspectives on the analysis results. S.Y.L. advised on the overall study workflow and results. This study was performed under the supervision and guidance of Y.K., H.Y., and S.Y., who hold responsibility for the final approval of this manuscript to be published.

## Methods

### 1 Collection of datasets and selection of samples

Two individual transcriptomic datasets from melanoma tumors were acquired and analyzed in this study: the Hugo et al. (Sequence Read Archive (SRA) Study: SRP070710; BioProject: PRJNA312948) and Riaz et al. (SRA Study: SRP094781; BioProject: PRJNA356761) datasets. Both datasets were generated from pretreated melanoma patients treated with anti-PD-1 immunotherapeutic agents (pembrolizumab or nivolumab) and were published elsewhere (Hugo et al., 2016; Riaz et al., 2017). The corresponding information on the patients’ mutational status and clinical classifications were collected from their supplementary materials.

In detail, paired-end whole transcriptome sequencing (WTS or RNA-seq) datasets in FASTQ format were downloaded from the SRA repository of the National Center for Biotechnology Information (NCBI). To maximize the distinction between groups, only the donors whose response to PD-1 inhibition was responding (complete response, CR and partial response, PR) or nonresponding (progressive disease, PD) were included in this study for the model establishment and development purposes.

Additionally, a WTS dataset generated from lung squamous carcinoma patients diagnosed at Samsung Medical Center (SMC) were used for external validation; all tissues collected in this process were obtained with the approval of the Institutional Review Board (SMC-2013-10-112 and SMC-2018-03-130).

### 2 Analysis of WTS datasets

All the target FASTQ sequences were aligned to the human (GRCh38.86 assembly) reference genome using the Spliced Transcript Alignment to a Reference (STAR) aligner with version 2.5.2B (Dobin et al., 2013). Then, the aligned reads were used to quantify gene expression and normalized into TPM units using the RNA-Seq by Expectation-Maximization (RSEM) software version 1.3.0 (Li and Dewey, 2011). As a result, we obtained the expression matrices from the two datasets and utilized them throughout this study.

### 3 LASSO regression

All subsequent numerical analyses were conducted using the *R* programming language, version 3.6.1, and its packages. To find an optimal gene list within the initial gene set, LASSO regression was performed for each dataset, with the anti-PD-1 response and the TPM values of 38 basal genes as dependent and independent variables, respectively. The LASSO regression has a variable selection effect by shrinking the estimators towards zero; hence, the variables in the regression model can be selected by removing the shrunken variables (i.e., zero-value estimator). After variable selection, the two lists of selected genes from each dataset were compared to identify commonly selected genes between the two independent datasets.

### 4 Application of the NMF method

We sought to identify underlying patterns between the two datasets regardless of their phenotype. For the regression-based approach, we applied the NMF method using the *R* packages *NMF*, version 0.22.0 (https://cran.r-project.org/package=NMF), and *NNLM*, version 0.4.3 (https://github.com/linxihui/NNLM). NMF factorizes an *N*×*P*matrix *A*, where *N* is the number of samples and *P* is the number of genes, into two matrices *W* and *H* with any rank *k* (*k < P*), where *WH ≈ A* (Lee and Seung, 1999). In many studies, NMF was hypothesized to decompose the gene expression matrix into multiple “metagene” axes. Thus, a metagene refers to a group of abstracted genes that contribute to the correlated expression. In the same manner, we employed NMF to extract major transcriptomic determinants, which are the genes contributing to each metagene referred to as matrix *W*. In our study, matrix *A* refers to a matrix of gene expression values for each patient. On the other hand, the decomposed matrices *W* and *H* consist of coefficients that show the relative contributions of each gene in each metagene and each metagene in each patient, respectively. The optimal value of a rank *k* was selected according to the cophenetic correlation coefficient (Sokal and Rohlf, 1962) from the rank survey analysis using matrix *A*. The empirical probability of the 4 metagenes from two independent datasets being identical was calculated with exhaustive enumeration.

### 5 Exhaustive search-based regression modeling

To find optimal predictors among the common metagene determinants, we performed exhaustive search-based regression subset selection in the final steps of model development using the R package *leaps*, v3.1 (https://cran.r-project.org/package=leaps). To compare stepwise predictive performance power, we demonstrated the exhaustive search for predictive modeling both with the *TDS* and without the *TDS*.

### 6 Conceptualization and calculation of the *TS* and *TDS*

In this paper, we hypothesized that immune checkpoint modulators have two effects: marginal effects and synergistic effects. While the marginal effect of an individual gene can be easily captured with the traditional regression framework, measuring synergistic effects is problematic. In this context, we introduced a simple concept named the *TS*. The *TS* is simply based on a one-to-all or exhaustive summation of coefficients of determination (*R*^2^) with the given gene set. Here, we derive the pairwise *R*^2^ as 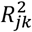 for the given expression matrix with *p* genes, which is defined as

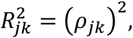

where *j* and *k* denote the *j*^th^ and *k*^th^ genes, respectively (*j,k* = 1, …,*p*). Here, we define two types of *TS*: gene set-wide and gene-wide *TS*. The former type of *TS* represents how much the genes in the gene set explain each other and is defined as

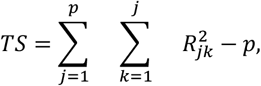

where *p* is the number of the genes, and 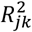 expresses an *R*^*2*^ value calculated by using two variables *j* and *k*. The latter type of *TS* represents how much a gene explains all other genes in the gene set; hence, it is defined as

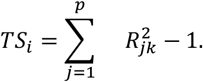

Furthermore, to quantify the contribution of each patient sample to the overall correlation pattern in the *TS*, we designed a subtraction-based calculation method for the *TDS*. The *TDS* value is calculated by subtracting *TS*_*(-n)*_ from *TS*_*Full*_, connoting the difference of the *TS* values when an individual variable (*n*) is eliminated.

Additionally, *abs_dTDS* were calculated by subtracting each *TDS* of pretreatment samples from that of on-treatment samples; these paired samples of 27 donors were included in the Riaz et al. dataset (Riaz et al., 2017).

### 7 Statistical analysis

To evaluate the statistical significance of *R*^2^ and the *TDS* between responders and nonresponders, we conducted Welch’s *t*-test on the group-wise vectors of *R*^2^ and the *TDS*, respectively.

To investigate the statistical properties of the *TS*, we computed its dataset-wide empirical distribution. For each of 10,000 replicates, the same number of genes (i.e., 38) included in the basal gene set were randomly selected, and the ratios of the responders’ *TS* to the nonresponders’ *TS* were calculated. The empirical distribution was derived by merging all those ratios, and its analytical distribution was estimated under the assumption of a normal distribution (**Figure S6A**).

To assess the significance of the gene-wise *TS* of the basal gene set, Wilcoxon rank-sum tests between the empirical distribution of the *TS* ratio and the empirical distributions of the target gene sets (the basal gene set and three Top38 gene sets) were performed. The significance of the individual *TS* ratio was assessed in both self-contained and competitive ways. We defined the empirical *p* value for those two approaches as its quantiles on the empirical distribution and its median-adjusted absolute distribution (see **Figures S6B-C** for both empirical distributions).

In the survival analysis, we derived and dichotomized the *TDS* using its median in a dataset-wise manner. We applied three Cox-PH models with dichotomized *TDS* against OS (in days) and vital status, which were suggested in the original articles: one dataset-wide model and two dataset-wise models. The LRT was used to derive *p* values. DEGs according to OS were also investigated in the three datasets, and the top 38 genes with the lowest *p* values were extracted for each dataset and compared across datasets.

### 8 Model performance evaluation

The predictive performance of the models mentioned in the text was evaluated using *k*-fold CV and vice versa prediction. We mainly used both 5- and 10-fold CV repeated 100 times and obtained mean test AUCs for each model. For relative comparison, we used various comparison models. The conventional models refer to models designed including approved targets. To minimize the bias within the 38 genes selected based on a thorough literature review with domain knowledge, we randomly selected 38 genes from the whole set of genes. In addition, we selected 38 top-ranked genes that were statistically marginal according to the p values from single-variable regression analyses in each dataset; the genes were selected from 17,286 genes included in the WTS dataset with an average TPM above 1. Vice versa prediction was performed by utilizing two independent melanoma datasets (the Hugo et al. (Hugo et al., 2016) and Riaz et al. (Riaz et al., 2017) datasets) as training and testing sets, respectively.

### 9 Estimation of immune cell fractions

We used a signature matrix of 22 functionally defined human immune subsets (LM22) derived from 547 marker genes to estimate fractions of immune cells by using CIBERSORTx (Newman et al., 2015; Newman et al., 2019) to implement a support vector regression (SVR)-based machine learning approach.

### 10 Alignment of protein sequences

To investigate sequence homology between the selected genes, we performed sequence alignment to compare protein sequences by employing BLASTp (protein-protein Basic Local Alignment Search Tool) (Altschul et al., 1990) available via the NCBI website. To run BLASTp, we entered either PD-1 or PD-L1 as the query sequence and each of the selected genes as the subject sequence. The output of the sequence alignment includes alignment scores, maximum scores, and total scores calculated between the query and subject sequences.

## Supplemental Information Titles and Legends

**Figure S1.**
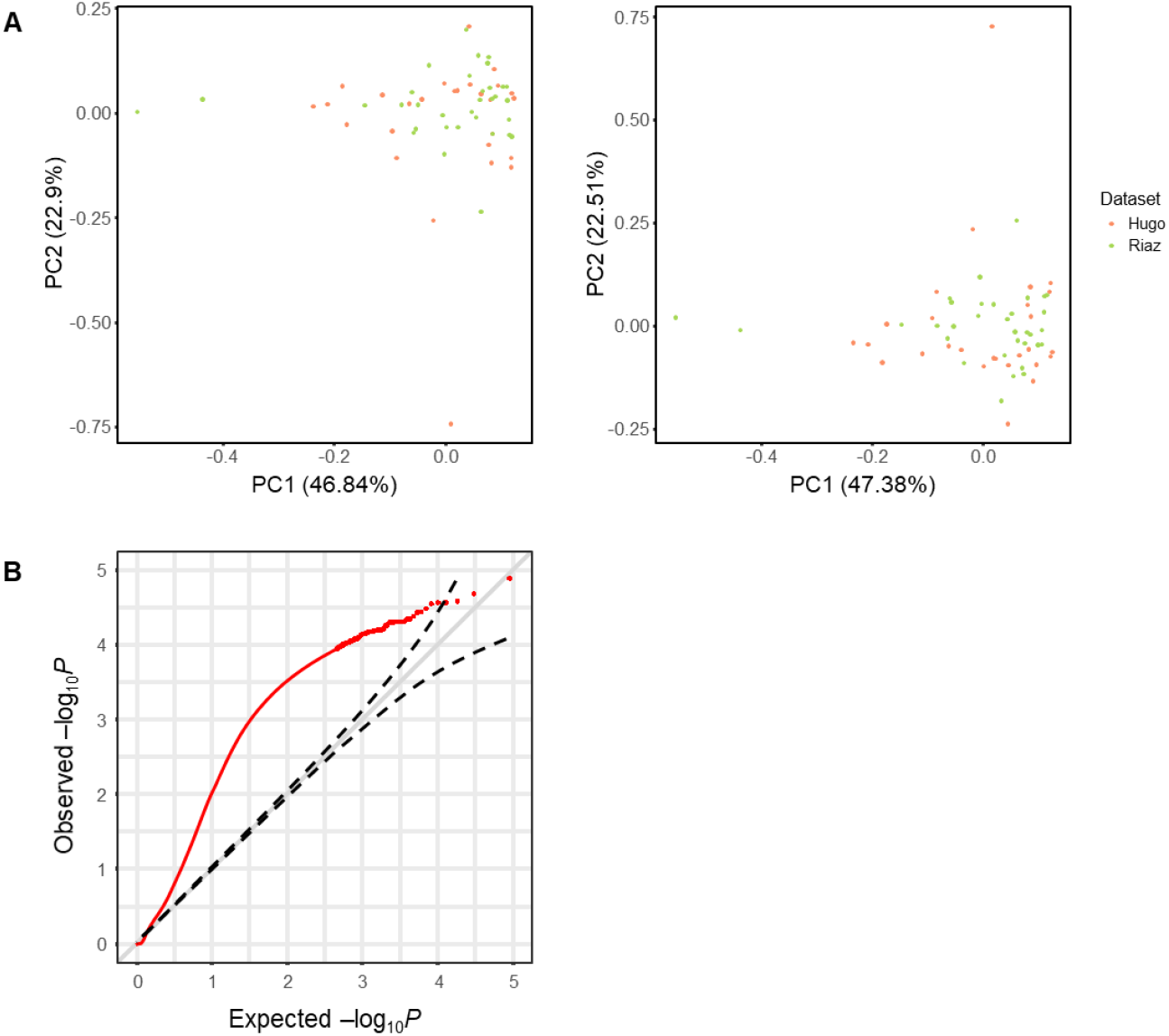
Heterogeneity of the two datasets used in the study. (A) PCA plots of the expression of the 38 immune checkpoint modulatory genes in the two datasets (Hugo et al. (Hugo et al., 2016) and Riaz et al. (Riaz et al., 2017)) before (**left**) and after (**right**) batch correction. (B) QQ plot of regression analysis based on the gene expression in each dataset. The plot demonstrates whether expression levels of the whole gene set are affected by the differences in each dataset (Hugo et al. (Hugo et al., 2016) and Riaz et al. (Riaz et al., 2017)).

**Figure S2.**
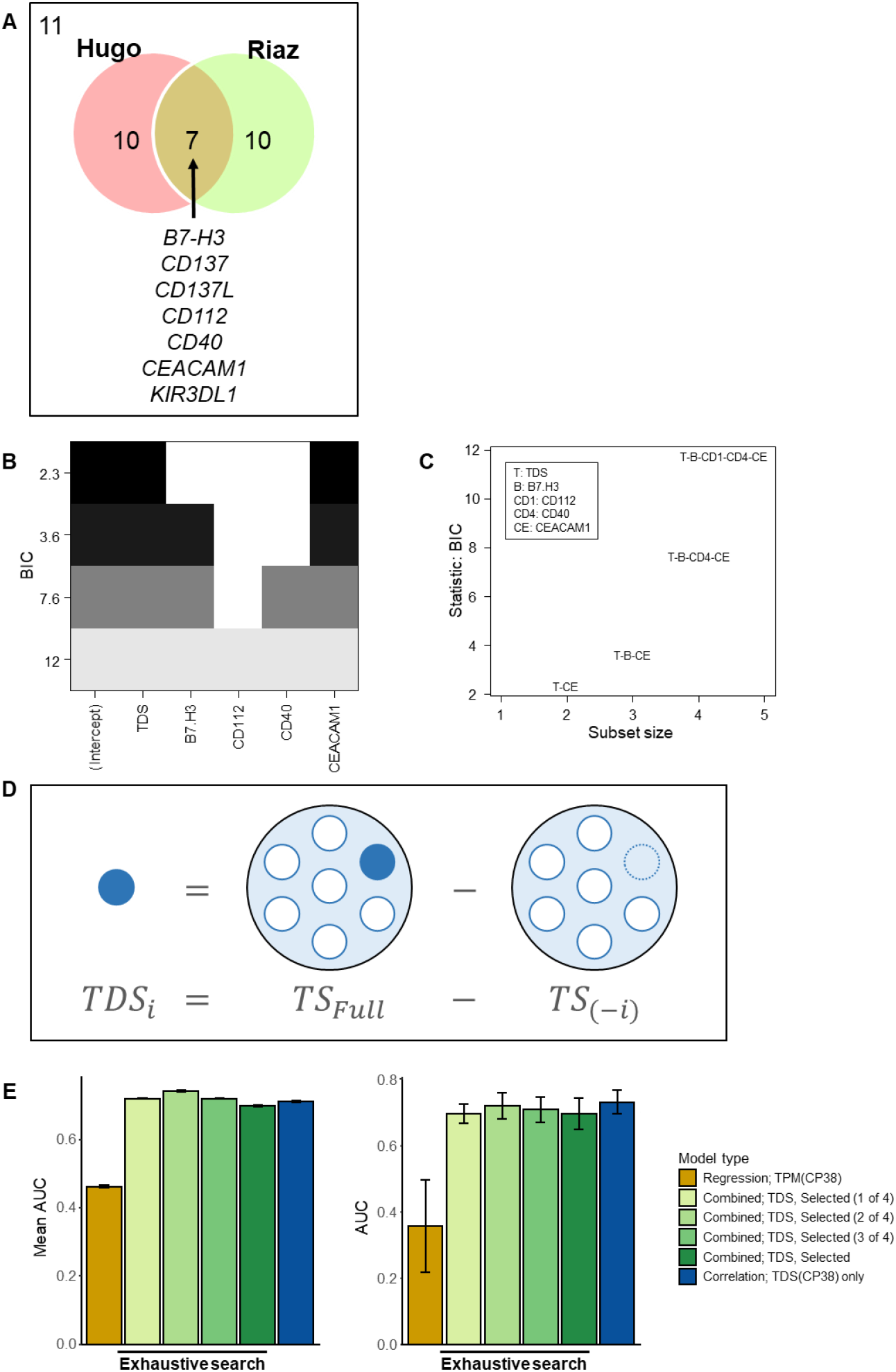
Regression-based approach: LASSO and the NMF method. (A) Venn diagram of selected genes from LASSO (Hugo vs. Riaz) showing the number of intersections and the differences in the added variables by forward selection between the two datasets. (B) Graphical table for exhaustive regression models (including the *TDS*). (C) Plot for exhaustive regression models (including the *TDS*). (D) AUCs from the 5-/10-fold CV and the vice versa prediction of exhaustive regression models (including the *TDS*). (E) This diagram illustrates the concept of calculating the *TDS* using *TS* values.

**Figure S3.**
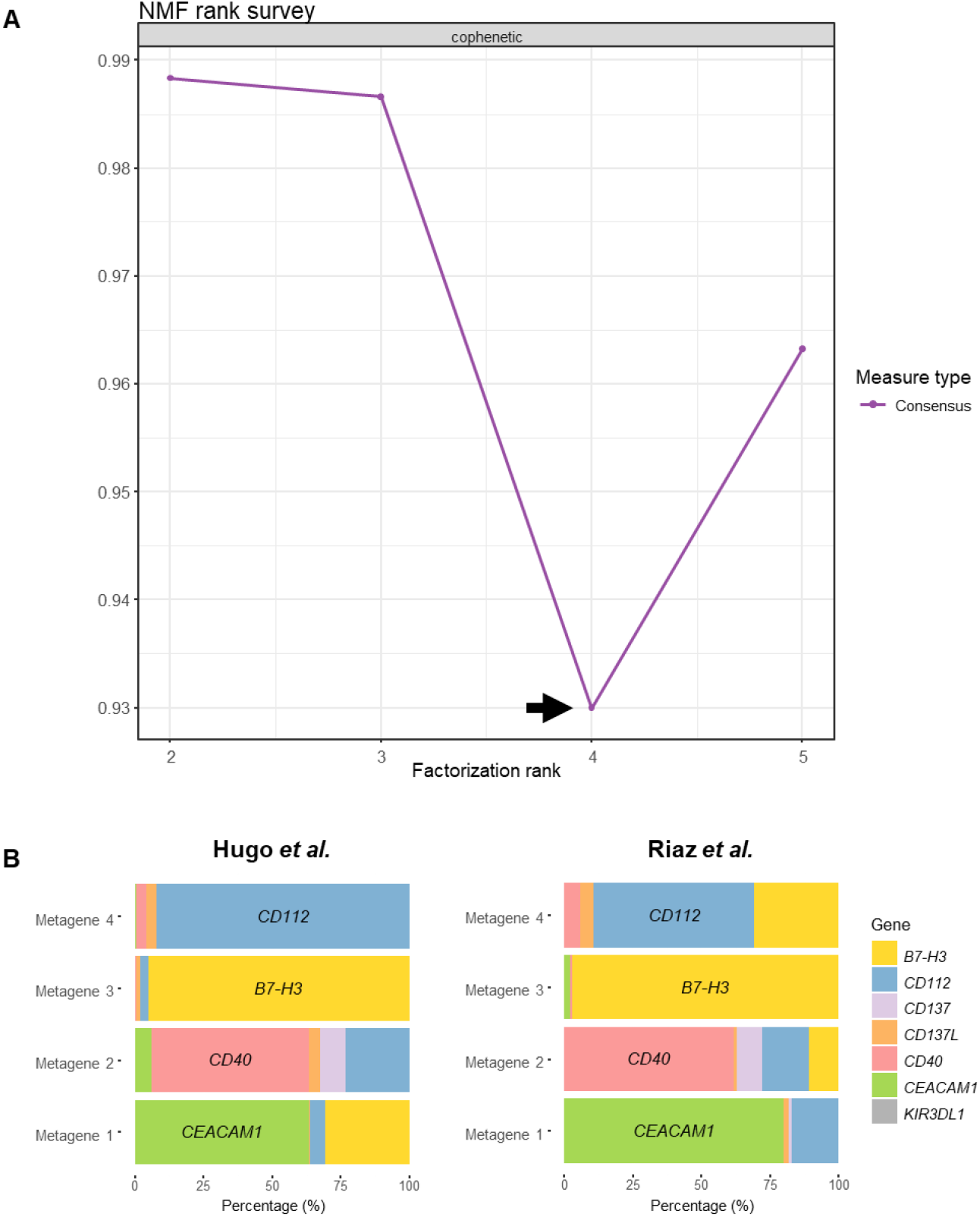
Supporting results from NMF. **(A) NMF rank survey**. The graph illustrates the cophenetic correlation coefficients estimated from the rank survey conducted to find an optimal rank for the NMF method. According to the lowest coefficient observed initially, the optimal rank was chosen as a value of 4. **(B) Metagene composition**. Each set of 4 stacked bar plots represents the composition of the 4 metagenes used in the study. Left: the metagene composition of the dataset of Hugo et al. (Hugo et al., 2016). Right: the metagene composition of the dataset of Riaz et al. (Riaz et al., 2017). The metagene composition of the merged dataset is depicted in **Figure 2B**.

**Figure S4.**
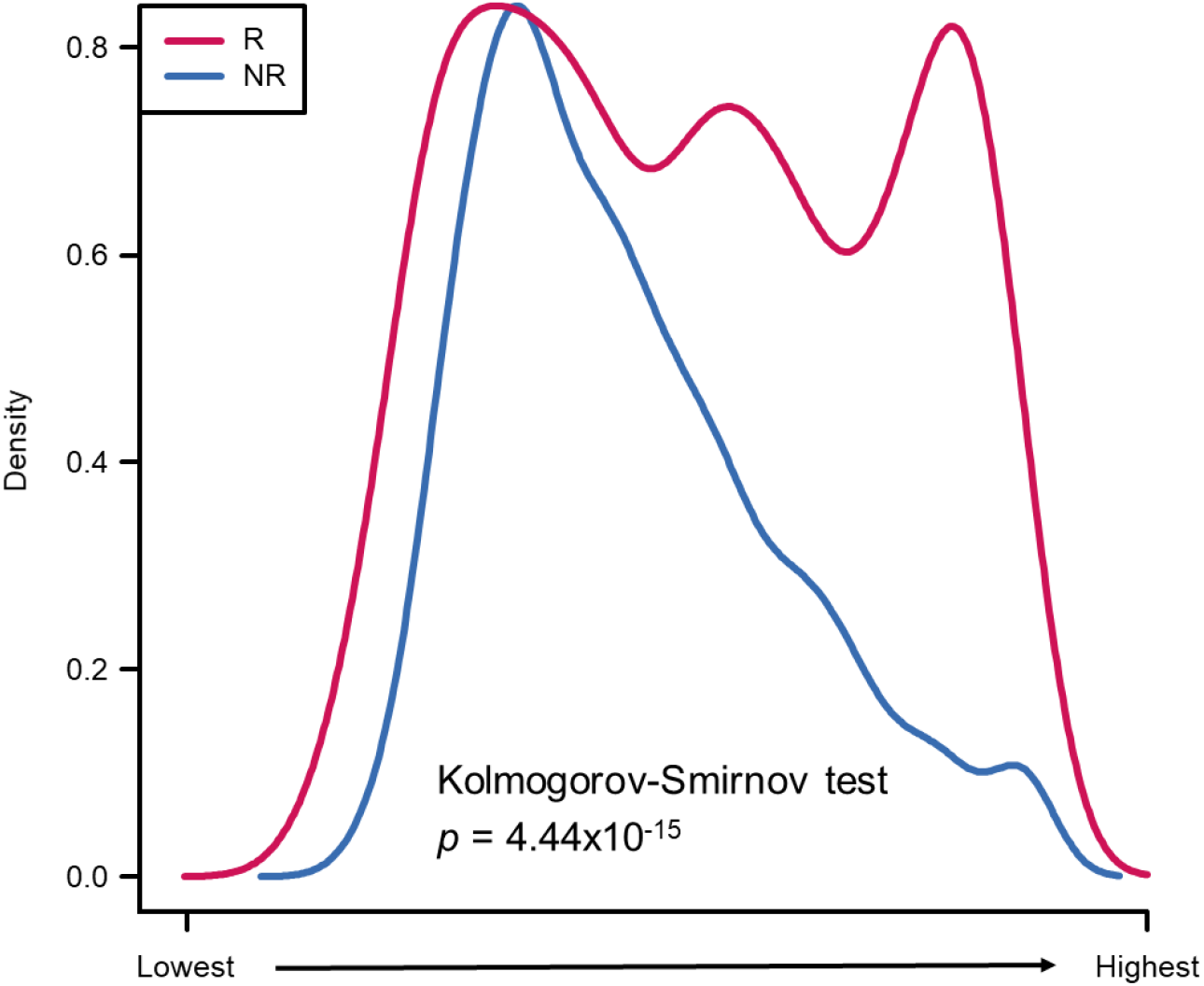
Kolmogorov-Smirnov (K-S) test for correlations of gene expression by response. This plot summarizes the overall distribution of correlations between the expression levels of the 38 immune checkpoint modulators by response group.

**Figure S5.**
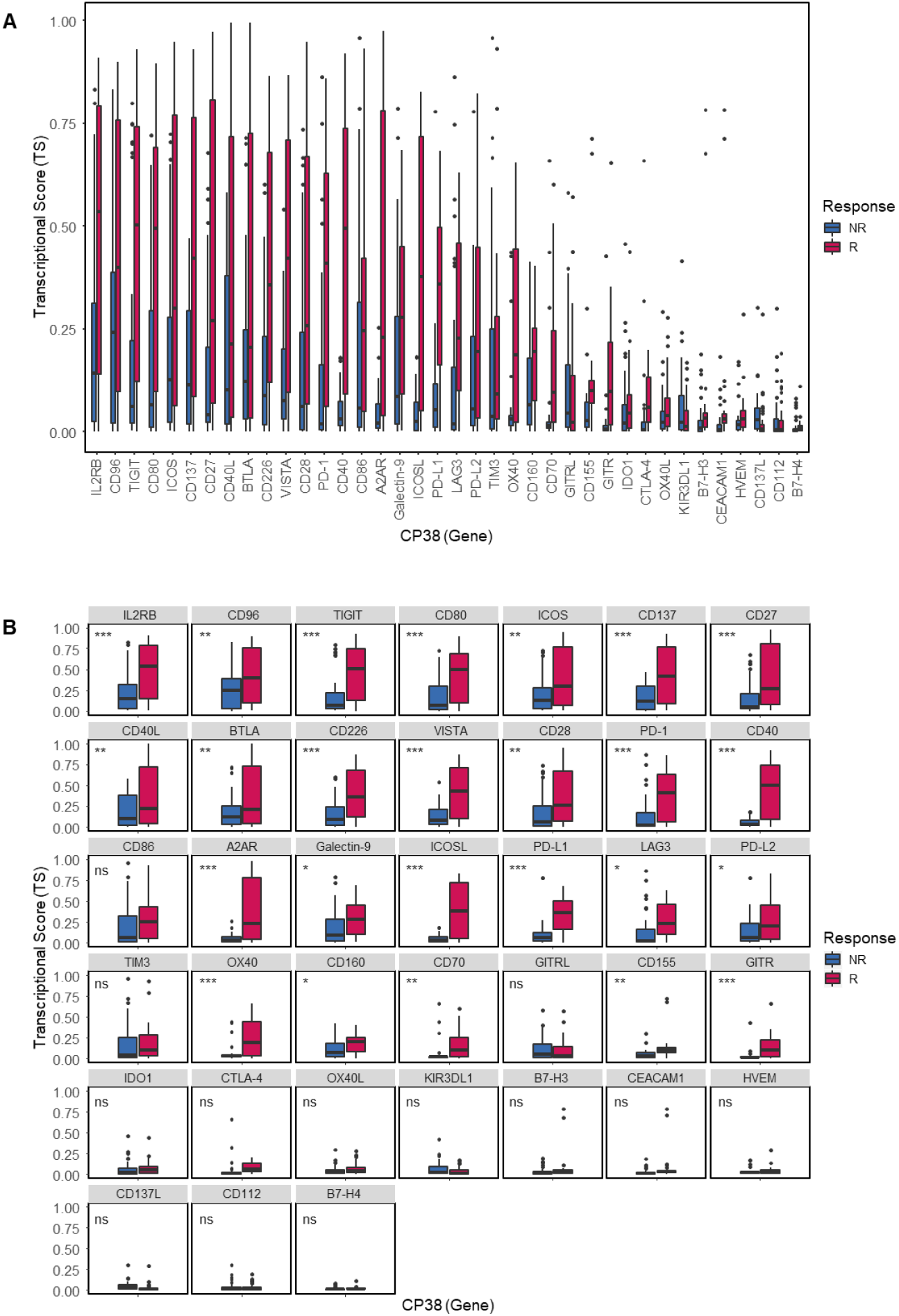
Coefficients of determination by response group. (A) Box plot of the *TS* calculated from the correlations among the expression levels of CP38 genes (in TPM) by response group. (B) Box plot of gene-wise *TS* calculated from the correlations among the expression levels of CP38 genes (in TPM) by response group. The statistical significance of the difference in expression of each gene between the two response groups is indicated with asterisks: *** (p < 0.001), ** (p ≤ 0.01), * (p ≤ 0.05), and ns (not significant; p > 0.05).

**Figure S6.**
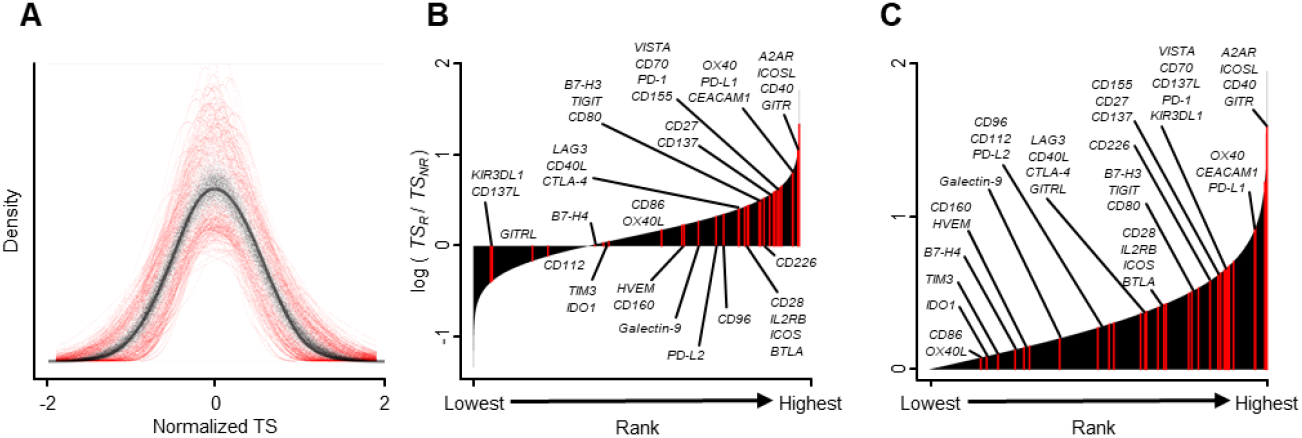
Properties of the *TS* and quantiles of the basal gene set. (A) The empirical distribution of 10,000 replicates of normalized *TS* from the combined dataset. (B) The empirical distribution of *TS* ratios by the responders’ *TS* over the nonresponders’ *TS* and quantiles of the basal gene set (red vertical lines) with the corresponding gene names (some are aggregated). (C) The empirical distribution of *TS* enrichment and quantiles of the basal gene set (red vertical lines) with the corresponding gene names (some are aggregated).

**Figure S7.**
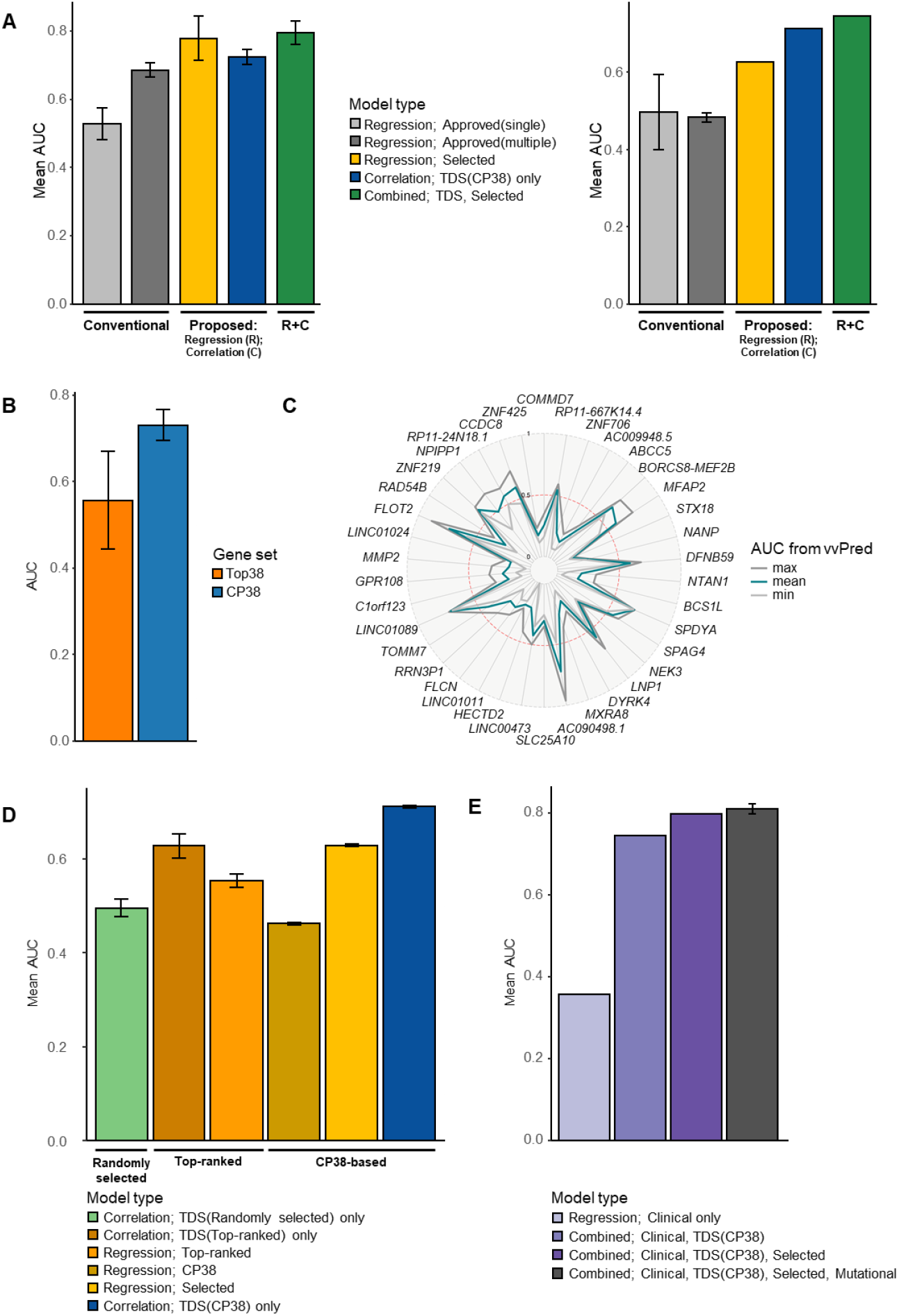
Additional performance evaluation of supporting models. (A) Mean AUCs from training sets and 5-fold CV in the merged datasets for melanoma. Each bar represents the predictive performance of the models. These models include either approved targets or proposed genes selected via our approaches. (B) AUCs from vice versa prediction using the Top38 and CP38 gene sets. Each Top38 gene set was derived from single-variable regression analysis of each dataset: H indicates Hugo et al. (Hugo et al., 2016), R indicates Riaz et al. (Riaz et al., 2017), and HR indicates the merged dataset containing the Hugo et al. and Riaz et al. datasets. (C) Radar plot of AUCs calculated from vice versa prediction using each of the 38 top-ranked genes in the HR dataset. The gene names are presented in descending order by AUC (clockwise). The red dotted line represents an AUC of 0.5; bold lines represent the lowest (min), mean, and highest (max) values of the prediction. (D) Summary of the model performance comparison. (E) Ten-fold cross-validated AUCs of models developed based on clinical features (i.e., M stage).

**Figure S8.**
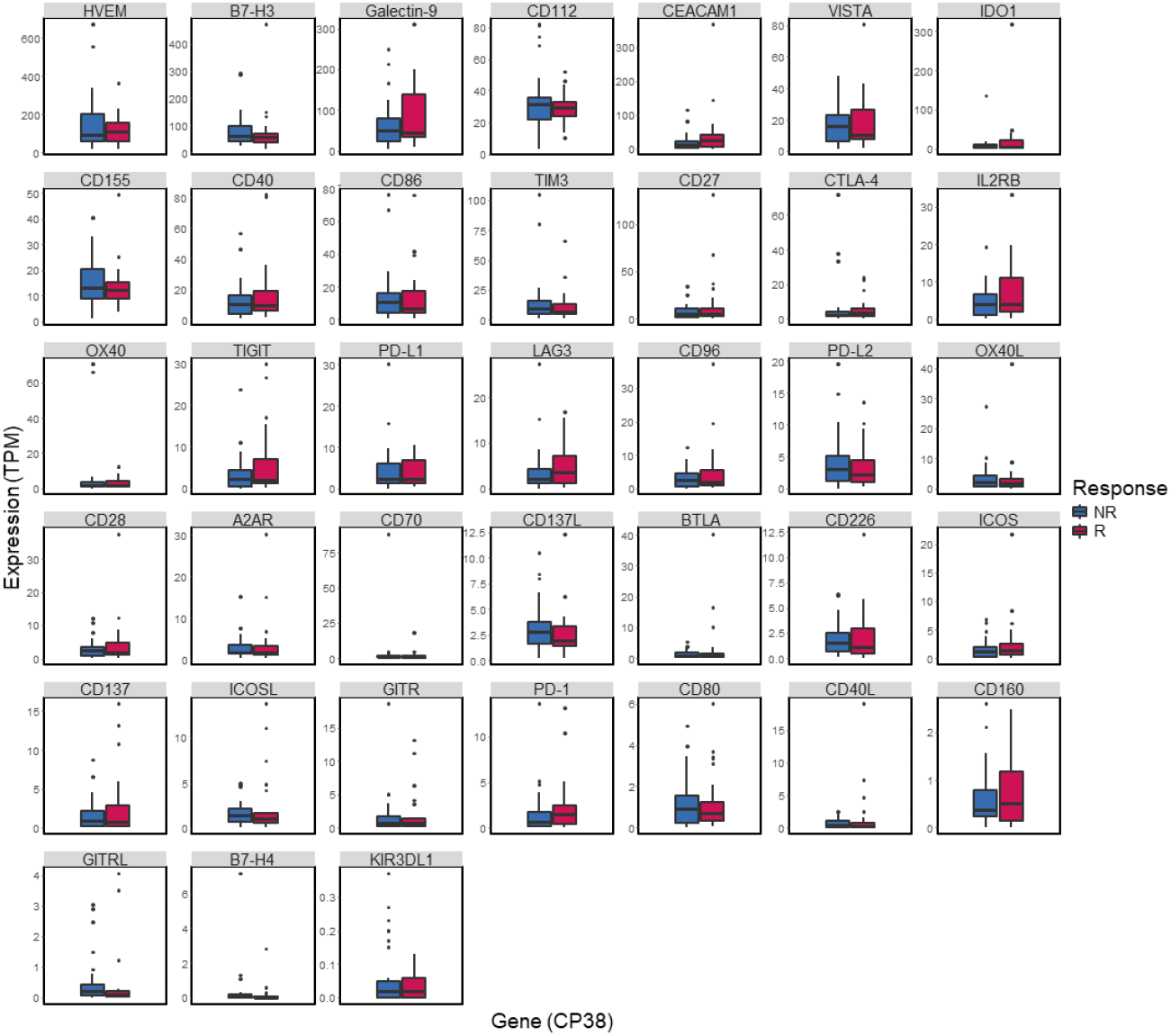
Basal expression of 38 immune checkpoint modulators in melanoma. Each bar plot represents the basal expression levels of 38 immune checkpoint modulators in TPM.

**Figure S9.**
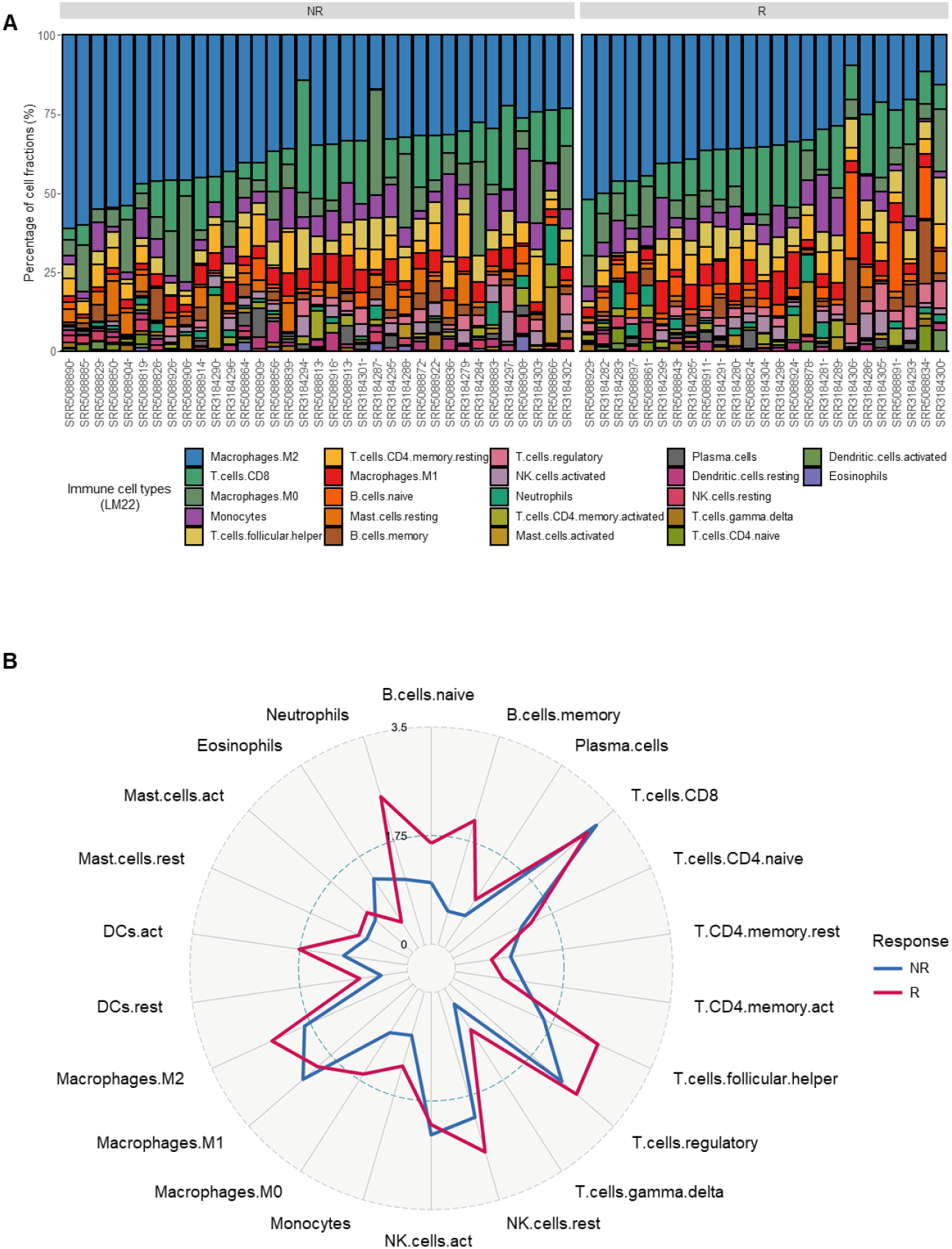
Distribution of immune cells by response (CIBERSORTx) (A) Stacked bar plot showing the fractions of 22 types of immune cells (LM22) estimated by CIBERSORTx; samples are sorted by the fractions of M2 macrophages, which were found to be the most frequent immune cell type in melanoma patients regardless of the anti-PD-1 response. Radar plot portraying the *TS* calculated by response group; immune cells are sorted by cell differentiation categories.

**Figure S10.**
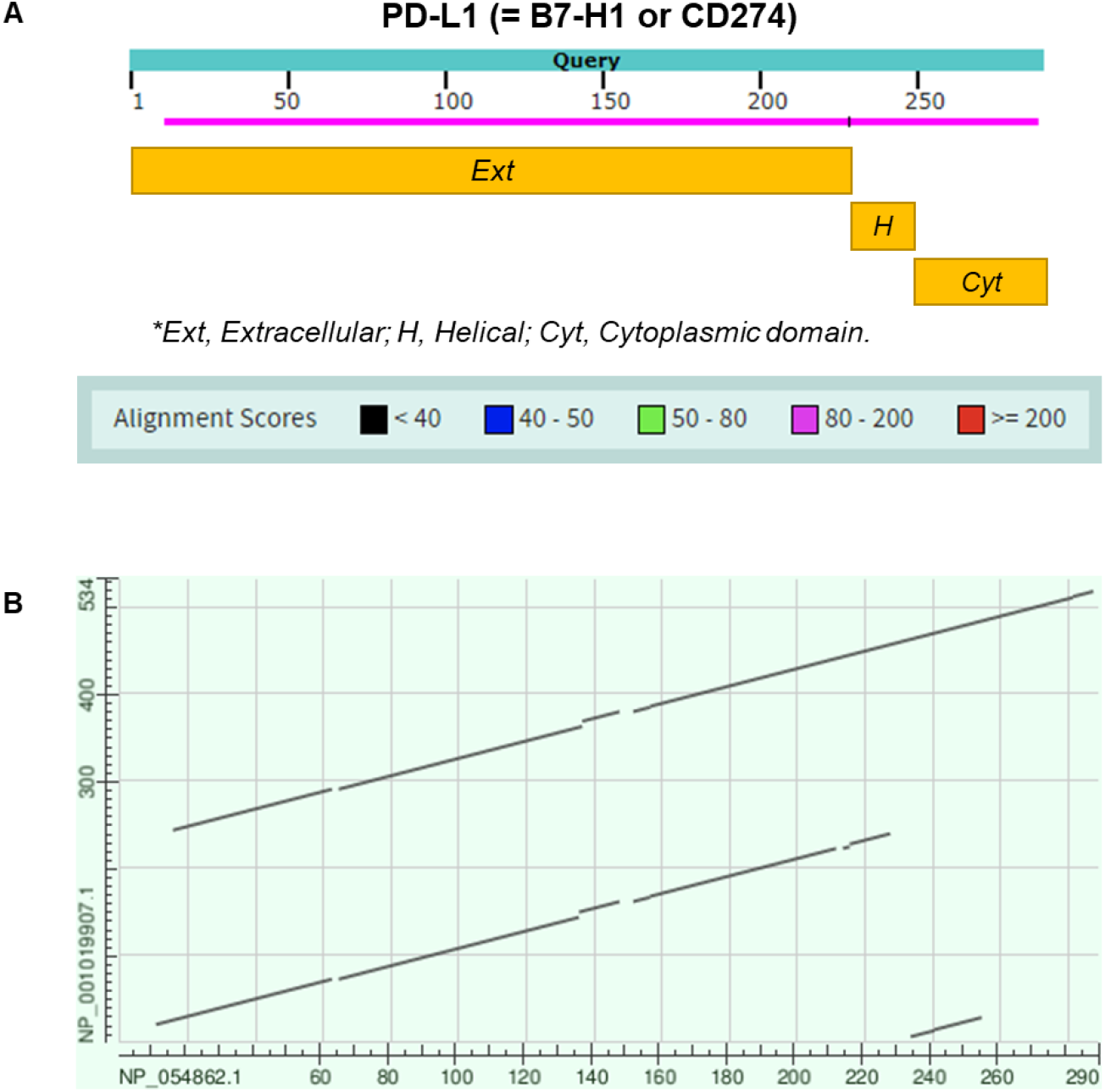
Sequence homology with PD-L1 (BLASTp) (A) Distribution of sequence alignment between B7-H3 and PD-L1 (B) Sequence alignment plot of B7-H3 and PD-L1

**Figure S11.**
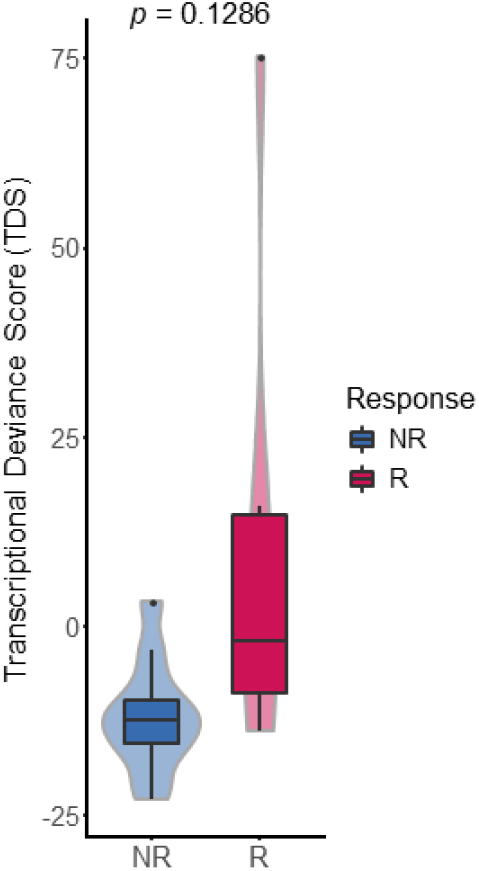
Distribution of the TDS in LUSC. This set of box plots shows the distribution of the *TDS* in LUSC. The statistical values were calculated by a one-sided Wilcoxon rank-sum exact test.

**Table S1.** List of 38 genes related to immune checkpoint modulators. This table shows the information on the 38 immune checkpoint modulatory genes used in the study; Ensembl gene IDs, HUGO Gene Nomenclature Committee (HGNC) symbols, gene names, and functional roles of each gene are included.

|  | Ensembl gene ID | HGNC symbol | Gene name | Functional role | Type |
| --- | --- | --- | --- | --- | --- |
| 1 | ENSG00000101017 | *CD40* | *CD40* | Stimulatory | Receptor |
| 2 | ENSG00000186891 | *TNFRSF18* | *GITR* | Stimulatory | Receptor |
| 3 | ENSG00000186827 | *TNFRSF4* | *OX40* | Stimulatory | Receptor |
| 4 | ENSG00000073008 | *PVR* | *CD155* | Stimulatory | Receptor |
| 5 | ENSG00000139193 | *CD27* | *CD27* | Stimulatory | Receptor |
| 6 | ENSG00000049249 | *TNFRSF9* | *CD137* | Stimulatory | Receptor |
| 7 | ENSG00000150637 | *CD226* | *CD226* | Stimulatory | Receptor |
| 8 | ENSG00000178562 | *CD28* | *CD28* | Stimulatory | Receptor |
| 9 | ENSG00000100385 | *IL2RB* | *IL2RB* | Stimulatory | Receptor |
| 10 | ENSG00000163600 | *ICOS* | *ICOS* | Stimulatory | Receptor |
| 11 | ENSG00000160223 | *ICOSLG* | *ICOSL* | Stimulatory | Ligand |
| 12 | ENSG00000125726 | *CD70* | *CD70* | Stimulatory | Ligand |
| 13 | ENSG00000121594 | *CD80* | *CD80* | Stimulatory | Ligand |
| 14 | ENSG00000102245 | *CD40LG* | *CD40L* | Stimulatory | Ligand |
| 15 | ENSG00000114013 | *CD86* | *CD86* | Stimulatory | Ligand |
| 16 | ENSG00000117586 | *TNFSF4* | *OX40L* | Stimulatory | Ligand |
| 17 | ENSG00000130202 | *NECTIN2* | *CD112* | Stimulatory | Ligand |
| 18 | ENSG00000120337 | *TNFSF18* | *GITRL* | Stimulatory | Ligand |
| 19 | ENSG00000125657 | *TNFSF9* | *CD137L* | Stimulatory | Ligand |
| 20 | ENSG00000128271 | *ADORA2A* | *A2AR* | Inhibitory | Receptor |
| 21 | ENSG00000107738 | *VSIR* | *VISTA* | Inhibitory | Receptor |
| 22 | ENSG00000188389 | *PDCD1* | *PD-1* | Inhibitory | Receptor |
| 23 | ENSG00000181847 | *TIGIT* | *TIGIT* | Inhibitory | Receptor |
| 24 | ENSG00000186265 | *BTLA* | *BTLA* | Inhibitory | Receptor |
| 25 | ENSG00000089692 | *LAG3* | *LAG3* | Inhibitory | Receptor |
| 26 | ENSG00000163599 | *CTLA4* | *CTLA-4* | Inhibitory | Receptor |
| 27 | ENSG00000153283 | *CD96* | *CD96* | Inhibitory | Receptor |
| 28 | ENSG00000117281 | *CD160* | *CD160* | Inhibitory | Receptor |
| 29 | ENSG00000135077 | *HAVCR2* | *TIM3* | Inhibitory | Receptor |
| 30 | ENSG00000167633 | *KIR3DL1* | *KIR3DL1* | Inhibitory | Receptor |
| 31 | ENSG00000079385 | *CEACAM1* | *CEACAM1* | Inhibitory | Receptor/Ligand |
| 32 | ENSG00000120217 | *CD274* | *PD-L1* | Inhibitory | Ligand |
| 33 | ENSG00000103855 | *CD276* | *B7-H3* | Inhibitory | Ligand |
| 34 | ENSG00000197646 | *PDCD1LG2* | *PD-L2* | Inhibitory | Ligand |
| 35 | ENSG00000168961 | *LGALS9* | *Galectin-9* | Inhibitory | Ligand |
| 36 | ENSG00000157873 | *TNFRSF14* | *HVEM* | Inhibitory | Ligand |
| 37 | ENSG00000131203 | *IDO1* | *IDO1* | Inhibitory | Ligand |
| 38 | ENSG00000134258 | *VTCN1* | *B7-H4* | Inhibitory | Ligand |

**Table S2.**
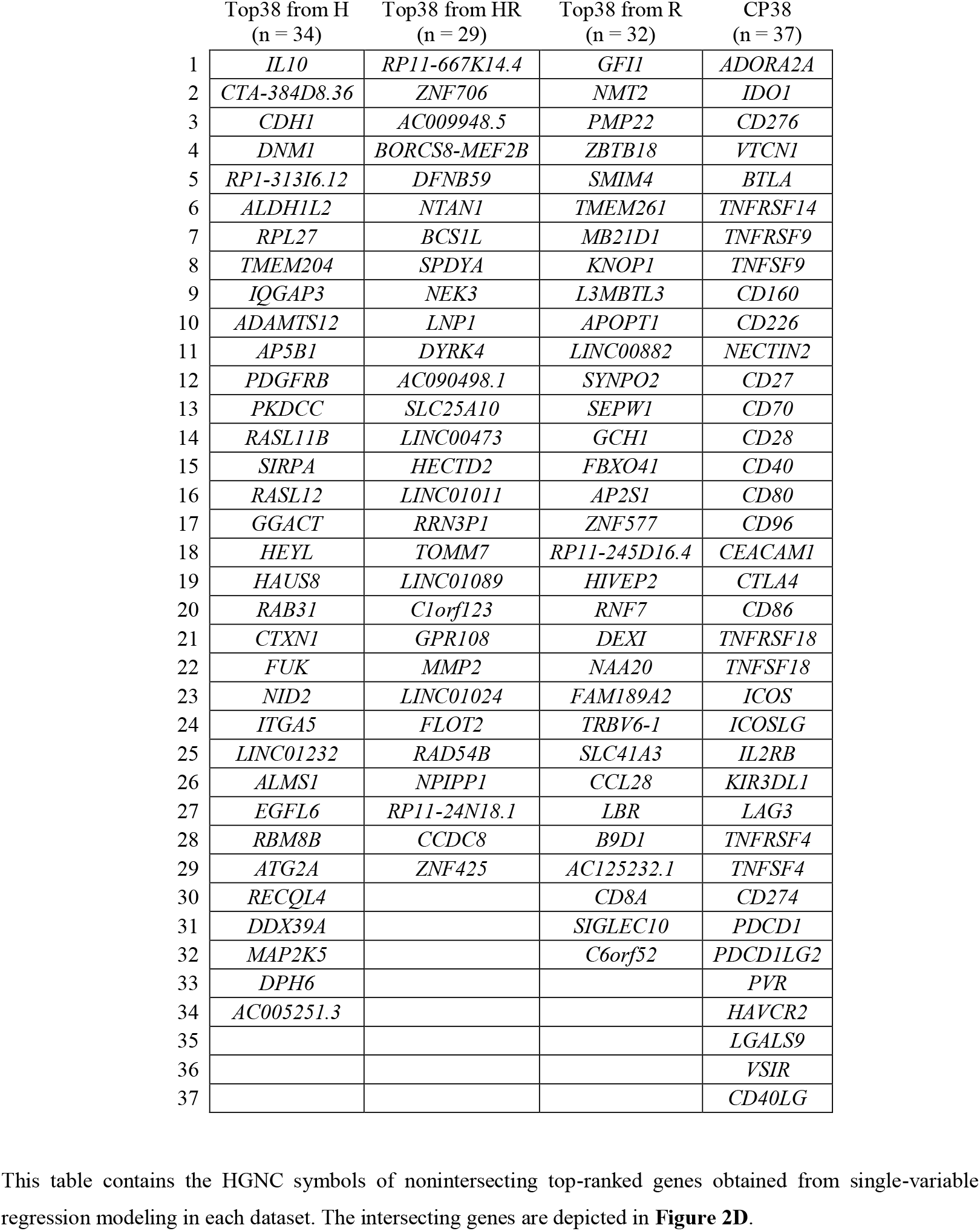
Top-ranked genes from single-variable regression modeling. This table contains the HGNC symbols of nonintersecting top-ranked genes obtained from single-variable regression modeling in each dataset. The intersecting genes are depicted in **Figure 2D**.

**Table S3.**
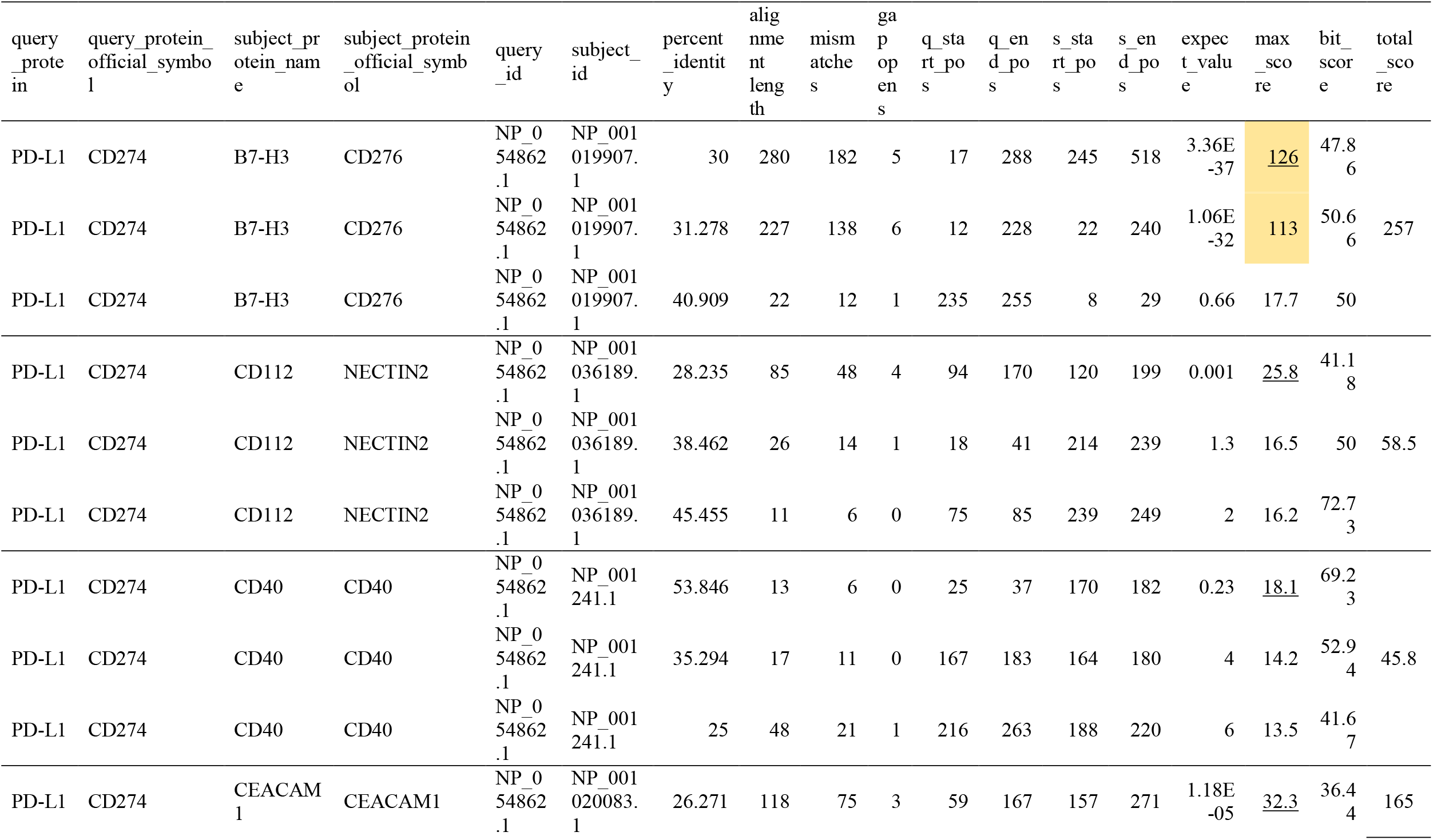
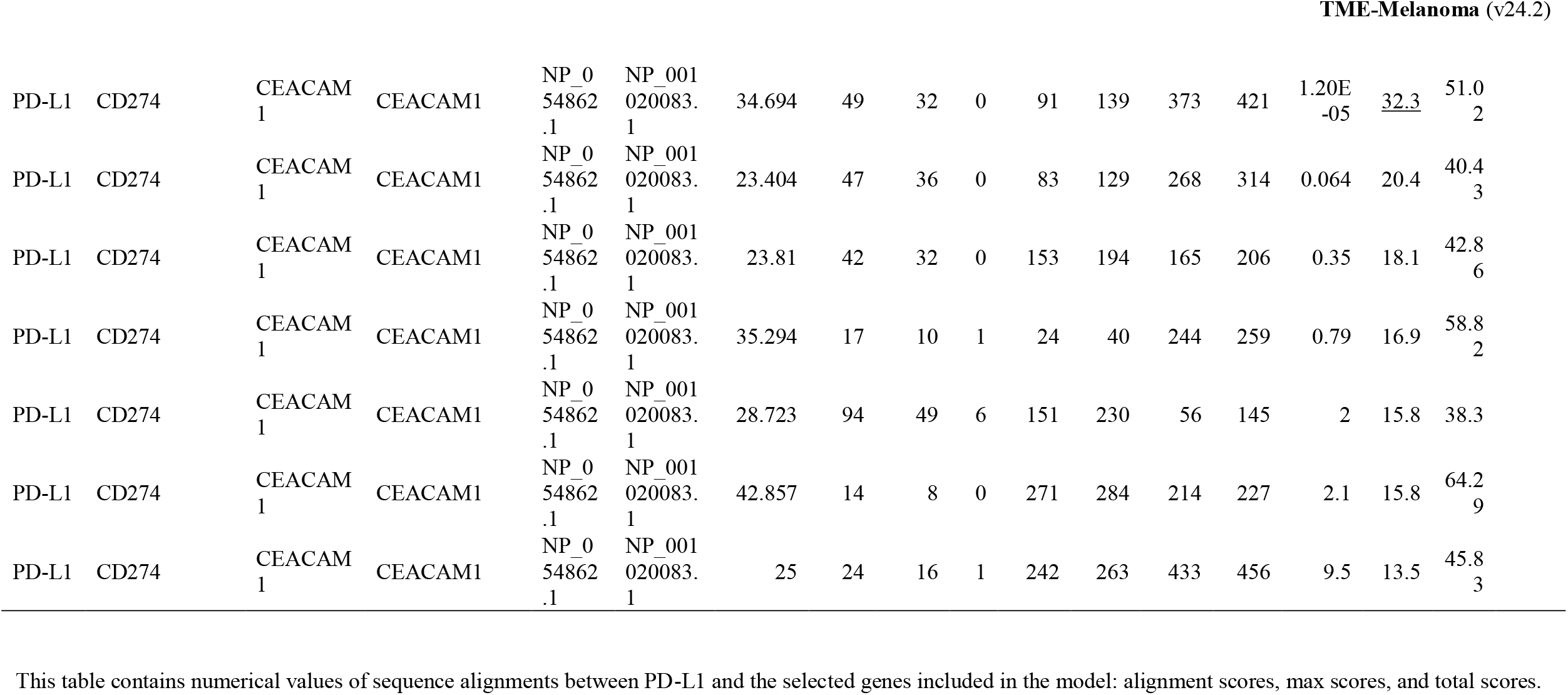
BLASTp results (alignment hit table for B7-H3 and PD-L1) This table contains numerical values of sequence alignments between PD-L1 and the selected genes included in the model: alignment scores, max scores, and total scores.

| query<br>_prote<br>in | query_protein_<br>official_symbo<br>l | subject_pr<br>oteins_nam<br>e | subject_protein<br>_official_symb<br>ol | query<br>_id | subject_<br>id | percent<br>_identit<br>y | alig<br>nme<br>nt<br>leng<br>th | mism<br>atche<br>s | ga<br>p<br>op<br>en<br>s | q_sta<br>rt_po<br>s | q_en<br>d_po<br>s | s_sta<br>rt_po<br>s | s_en<br>d_po<br>s | expec<br>t_valu<br>e | max<br>_sco<br>re | bit_<br>scor<br>e | total<br>_sco<br>re |
| --- | --- | --- | --- | --- | --- | --- | --- | --- | --- | --- | --- | --- | --- | --- | --- | --- | --- |
| PD-L1 | CD274 | B7-H3 | CD276 | NP_054862.1 | NP_001019907.1 | 30 | 280 | 182 | 5 | 17 | 288 | 245 | 518 | 3.36E-37 | <u>126</u> | 47.86 |  |
| PD-L1 | CD274 | B7-H3 | CD276 | NP_054862.1 | NP_001019907.1 | 31.278 | 227 | 138 | 6 | 12 | 228 | 22 | 240 | 1.06E-32 | <u>113</u> | 50.66 | 257 |
| PD-L1 | CD274 | B7-H3 | CD276 | NP_054862.1 | NP_001019907.1 | 40.909 | 22 | 12 | 1 | 235 | 255 | 8 | 29 | 0.66 | 17.7 | 50 |  |
| PD-L1 | CD274 | CD112 | NECTIN2 | NP_054862.1 | NP_001036189.1 | 28.235 | 85 | 48 | 4 | 94 | 170 | 120 | 199 | 0.001 | <u>25.8</u> | 41.18 |  |
| PD-L1 | CD274 | CD112 | NECTIN2 | NP_054862.1 | NP_001036189.1 | 38.462 | 26 | 14 | 1 | 18 | 41 | 214 | 239 | 1.3 | 16.5 | 50 | 58.5 |
| PD-L1 | CD274 | CD112 | NECTIN2 | NP_054862.1 | NP_001036189.1 | 45.455 | 11 | 6 | 0 | 75 | 85 | 239 | 249 | 2 | 16.2 | 72.73 |  |
| PD-L1 | CD274 | CD40 | CD40 | NP_054862.1 | NP_001241.1 | 53.846 | 13 | 6 | 0 | 25 | 37 | 170 | 182 | 0.23 | <u>18.1</u> | 69.23 |  |
| PD-L1 | CD274 | CD40 | CD40 | NP_054862.1 | NP_001241.1 | 35.294 | 17 | 11 | 0 | 167 | 183 | 164 | 180 | 4 | 14.2 | 52.94 | 45.8 |
| PD-L1 | CD274 | CD40 | CD40 | NP_054862.1 | NP_001241.1 | 25 | 48 | 21 | 1 | 216 | 263 | 188 | 220 | 6 | 13.5 | 41.67 |  |
| PD-L1 | CD274 | CEACAM1 | CEACAM1 | NP_054862.1 | NP_001020083.1 | 26.271 | 118 | 75 | 3 | 59 | 167 | 157 | 271 | 1.18E-05 | <u>32.3</u> | 36.44 | 165 |

|  |  |  |  |  |  |  |  |  |  |  |  |  |  |  |  |  |
| --- | --- | --- | --- | --- | --- | --- | --- | --- | --- | --- | --- | --- | --- | --- | --- | --- |
| PD-L1 | CD274 | CEACAM1 | CEACAM1 | NP_054862.1 | NP_001020083.1 | 34.694 | 49 | 32 | 0 | 91 | 139 | 373 | 421 | 1.20E-05 | <u>32.3</u> | 51.02 |
| PD-L1 | CD274 | CEACAM1 | CEACAM1 | NP_054862.1 | NP_001020083.1 | 23.404 | 47 | 36 | 0 | 83 | 129 | 268 | 314 | 0.064 | 20.4 | 40.43 |
| PD-L1 | CD274 | CEACAM1 | CEACAM1 | NP_054862.1 | NP_001020083.1 | 23.81 | 42 | 32 | 0 | 153 | 194 | 165 | 206 | 0.35 | 18.1 | 42.86 |
| PD-L1 | CD274 | CEACAM1 | CEACAM1 | NP_054862.1 | NP_001020083.1 | 35.294 | 17 | 10 | 1 | 24 | 40 | 244 | 259 | 0.79 | 16.9 | 58.82 |
| PD-L1 | CD274 | CEACAM1 | CEACAM1 | NP_054862.1 | NP_001020083.1 | 28.723 | 94 | 49 | 6 | 151 | 230 | 56 | 145 | 2 | 15.8 | 38.3 |
| PD-L1 | CD274 | CEACAM1 | CEACAM1 | NP_054862.1 | NP_001020083.1 | 42.857 | 14 | 8 | 0 | 271 | 284 | 214 | 227 | 2.1 | 15.8 | 64.29 |
| PD-L1 | CD274 | CEACAM1 | CEACAM1 | NP_054862.1 | NP_001020083.1 | 25 | 24 | 16 | 1 | 242 | 263 | 433 | 456 | 9.5 | 13.5 | 45.83 |

## References

Alexandrov, L. B., Nik-Zainal, S., Wedge, D. C., Aparicio, S. A., Behjati, S., Biankin, A. V., Bignell, G. R., Bolli, N., Borg, A., Borresen-Dale, A. L., et al. (2013). Signatures of mutational processes in human cancer. Nature 500, 415–421.

Altschul, S. F., Gish, W., Miller, W., Myers, E. W., and Lipman, D. J. (1990). Basic local alignment search tool. J Mol Biol 215, 403–410.

Ancevski Hunter, K., Socinski, M. A., and Villaruz, L. C. (2018). PD-L1 Testing in Guiding Patient Selection for PD-1/PD-L1 Inhibitor Therapy in Lung Cancer. Mol Diagn Ther 22, 1–10.

Andrews, L. P., Yano, H., and Vignali, D. A. A. (2019). Inhibitory receptors and ligands beyond PD-1, PD-L1 and CTLA-4: breakthroughs or backups. Nat Immunol 20, 1425–1434.

Auslander, N., Lee, J. S., and Ruppin, E. (2019). Reply to: ‘IMPRES does not reproducibly predict response to immune checkpoint blockade therapy in metastatic melanoma’. Nat Med 25, 1836–1838.

Auslander, N., Zhang, G., Lee, J. S., Frederick, D. T., Miao, B., Moll, T., Tian, T., Wei, Z., Madan, S., Sullivan, R. J., et al. (2018). Publisher Correction: Robust prediction of response to immune checkpoint blockade therapy in metastatic melanoma. Nat Med 24, 1942.

Beavis, P. A., Milenkovski, N., Henderson, M. A., John, L. B., Allard, B., Loi, S., Kershaw, M. H., Stagg, J., and Darcy, P. K. (2015). Adenosine Receptor 2A Blockade Increases the Efficacy of Anti-PD-1 through Enhanced Antitumor T-cell Responses. Cancer Immunol Res 3, 506–517.

Cabrita, R., Lauss, M., Sanna, A., Donia, M., Skaarup Larsen, M., Mitra, S., Johansson, I., Phung, B., Harbst, K., Vallon-Christersson, J., et al. (2020). Tertiary lymphoid structures improve immunotherapy and survival in melanoma. Nature.

Cai, D., Li, J., Liu, D., Hong, S., Qiao, Q., Sun, Q., Li, P., Lyu, N., Sun, T., Xie, S., et al. (2019). Tumor-expressed B7-H3 mediates the inhibition of antitumor T-cell functions in ovarian cancer insensitive to PD-1 blockade therapy. Cell Mol Immunol.

Carter, J. A., Gilbo, P., and Atwal, G. S. (2019). IMPRES does not reproducibly predict response to immune checkpoint blockade therapy in metastatic melanoma. Nat Med 25, 1833–1835.

Castellanos, J. R., Purvis, I. J., Labak, C. M., Guda, M. R., Tsung, A. J., Velpula, K. K., and Asuthkar, S. (2017). B7-H3 role in the immune landscape of cancer. Am J Clin Exp Immunol 6, 66–75.

Cekic, C., and Linden, J. (2014). Adenosine A2A receptors intrinsically regulate CD8+ T cells in the tumor microenvironment. Cancer Res 74, 7239–7249.

Chapoval, A. I., Ni, J., Lau, J. S., Wilcox, R. A., Flies, D. B., Liu, D., Dong, H., Sica, G. L., Zhu, G., Tamada, K., and Chen, L. (2001). B7-H3: a costimulatory molecule for T cell activation and IFN-gamma production. Nat Immunol 2, 269–274.

Charoentong, P., Finotello, F., Angelova, M., Mayer, C., Efremova, M., Rieder, D., Hackl, H., and Trajanoski, Z. (2017). Pan-cancer Immunogenomic Analyses Reveal Genotype-Immunophenotype Relationships and Predictors of Response to Checkpoint Blockade. Cell Rep 18, 248–262.

Chretien, S., Zerdes, I., Bergh, J., Matikas, A., and Foukakis, T. (2019). Beyond PD-1/PD-L1 Inhibition: What the Future Holds for Breast Cancer Immunotherapy. Cancers (Basel) 11.

Danaher, P., Warren, S., Lu, R., Samayoa, J., Sullivan, A., Pekker, I., Wallden, B., Marincola, F. M., and Cesano, A. (2018). Pan-cancer adaptive immune resistance as defined by the Tumor Inflammation Signature (TIS): results from The Cancer Genome Atlas (TCGA). J Immunother Cancer 6, 63.

Dankner, M., Gray-Owen, S. D., Huang, Y. H., Blumberg, R. S., and Beauchemin, N. (2017). CEACAM1 as a multi-purpose target for cancer immunotherapy. Oncoimmunology 6, e1328336.

Darvin, P., Toor, S. M., Sasidharan Nair, V., and Elkord, E. (2018). Immune checkpoint inhibitors: recent progress and potential biomarkers. Exp Mol Med 50, 165.

Davis, A. A., and Patel, V. G. (2019). The role of PD-L1 expression as a predictive biomarker: an analysis of all US Food and Drug Administration (FDA) approvals of immune checkpoint inhibitors. J Immunother Cancer 7, 278.

De Silva, N. S., and Klein, U. (2015). Dynamics of B cells in germinal centres. Nat Rev Immunol 15, 137–148.

Dobin, A., Davis, C. A., Schlesinger, F., Drenkow, J., Zaleski, C., Jha, S., Batut, P., Chaisson, M., and Gingeras, T. R. (2013). STAR: ultrafast universal RNA-seq aligner. Bioinformatics 29, 15–21.

Dong, H., Strome, S. E., Salomao, D. R., Tamura, H., Hirano, F., Flies, D. B., Roche, P. C., Lu, J., Zhu, G., Tamada, K., et al. (2002). Tumor-associated B7-H1 promotes T-cell apoptosis: a potential mechanism of immune evasion. Nat Med 8, 793–800.

Flem-Karlsen, K., Fodstad, Y., and Nunes-Xavier, C. E. (2019). B7-H3 immune checkpoint protein in human cancer. Curr Med Chem.

Fridman, W. H., Pages, F., Sautes-Fridman, C., and Galon, J. (2012). The immune contexture in human tumours: impact on clinical outcome. Nat Rev Cancer 12, 298–306.

Galon, J., and Bruni, D. (2019). Approaches to treat immune hot, altered and cold tumours with combination immunotherapies. Nat Rev Drug Discov 18, 197–218.

Galon, J., Fox, B. A., Bifulco, C. B., Masucci, G., Rau, T., Botti, G., Marincola, F. M., Ciliberto, G., Pages, F., Ascierto, P. A., and Capone, M. (2016). Immunoscore and Immunoprofiling in cancer: an update from the melanoma and immunotherapy bridge 2015. J Transl Med 14, 273.

Garon, E. B., Rizvi, N. A., Hui, R., Leighl, N., Balmanoukian, A. S., Eder, J. P., Patnaik, A., Aggarwal, C., Gubens, M., Horn, L., et al. (2015). Pembrolizumab for the treatment of non-small-cell lung cancer. N Engl J Med 372, 2018–2028.

Geukes Foppen, M. H., Donia, M., Svane, I. M., and Haanen, J. B. (2015). Tumor-infiltrating lymphocytes for the treatment of metastatic cancer. Mol Oncol 9, 1918–1935.

Goodman, A. M., Kato, S., Bazhenova, L., Patel, S. P., Frampton, G. M., Miller, V., Stephens, P. J., Daniels, G. A., and Kurzrock, R. (2017). Tumor Mutational Burden as an Independent Predictor of Response to Immunotherapy in Diverse Cancers. Mol Cancer Ther 16, 2598–2608.

Guo, F. F., and Cui, J. W. (2019). The Role of Tumor-Infiltrating B Cells in Tumor Immunity. J Oncol 2019, 2592419.

Hayward, N. K., Wilmott, J. S., Waddell, N., Johansson, P. A., Field, M. A., Nones, K., Patch, A. M., Kakavand, H., Alexandrov, L. B., Burke, H., et al. (2017). Whole-genome landscapes of major melanoma subtypes. Nature 545, 175–180.

Hugo, W., Zaretsky, J. M., Sun, L., Song, C., Moreno, B. H., Hu-Lieskovan, S., Berent-Maoz, B., Pang, J., Chmielowski, B., Cherry, G., et al. (2016). Genomic and Transcriptomic Features of Response to Anti-PD-1 Therapy in Metastatic Melanoma. Cell 165, 35–44.

Keenan, T. E., Burke, K. P., and Van Allen, E. M. (2019). Genomic correlates of response to immune checkpoint blockade. Nat Med 25, 389–402.

Kim, J., Cho, J., Lee, M. H., and Lim, J. H. (2018). Relative Efficacy of Checkpoint Inhibitors for Advanced NSCLC According to Programmed Death-Ligand-1 Expression: A Systematic Review and Network Meta-Analysis. Sci Rep 8, 11738.

Kitano, S., Nakayama, T., and Yamashita, M. (2018). Biomarkers for Immune Checkpoint Inhibitors in Melanoma. Front Oncol 8, 270.

Kornbluth, R. S., Stempniak, M., and Stone, G. W. (2012). Design of CD40 agonists and their use in growing B cells for cancer immunotherapy. Int Rev Immunol 31, 279–288.

Lee, D. D., and Seung, H. S. (1999). Learning the parts of objects by non-negative matrix factorization. Nature 401, 788–791.

Lee, Y. H., Martin-Orozco, N., Zheng, P., Li, J., Zhang, P., Tan, H., Park, H. J., Jeong, M., Chang, S. H., Kim, B. S., et al. (2017). Inhibition of the B7-H3 immune checkpoint limits tumor growth by enhancing cytotoxic lymphocyte function. Cell Res 27, 1034–1045.

Leone, R. D., Sun, I. M., Oh, M. H., Sun, I. H., Wen, J., Englert, J., and Powell, J. D. (2018). Inhibition of the adenosine A2a receptor modulates expression of T cell coinhibitory receptors and improves effector function for enhanced checkpoint blockade and ACT in murine cancer models. Cancer Immunol Immunother 67, 1271–1284.

Li, B., and Dewey, C. N. (2011). RSEM: accurate transcript quantification from RNA-Seq data with or without a reference genome. BMC Bioinformatics 12, 323.

Lines, J. L., Sempere, L. F., Broughton, T., Wang, L., and Noelle, R. (2014). VISTA is a novel broad-spectrum negative checkpoint regulator for cancer immunotherapy. Cancer Immunol Res 2, 510–517.

Liu, D., Schilling, B., Liu, D., Sucker, A., Livingstone, E., Jerby-Amon, L., Zimmer, L., Gutzmer, R., Satzger, I., Loquai, C., et al. (2019). Integrative molecular and clinical modeling of clinical outcomes to PD1 blockade in patients with metastatic melanoma. Nat Med 25, 1916–1927.

Luke, J. J., and Ascierto, P. A. (2020). Biology confirmed but biomarkers elusive in melanoma immunotherapy. Nat Rev Clin Oncol 17, 198–199.

Luke, J. J., Flaherty, K. T., Ribas, A., and Long, G. V. (2017). Targeted agents and immunotherapies: optimizing outcomes in melanoma. Nat Rev Clin Oncol 14, 463–482.

Martin-Orozco, N., Li, Y., Wang, Y., Liu, S., Hwu, P., Liu, Y. J., Dong, C., and Radvanyi, L. (2010). Melanoma cells express ICOS ligand to promote the activation and expansion of T-regulatory cells. Cancer Res 70, 9581–9590.

McLaughlin, J., Han, G., Schalper, K. A., Carvajal-Hausdorf, D., Pelekanou, V., Rehman, J., Velcheti, V., Herbst, R., LoRusso, P., and Rimm, D. L. (2016). Quantitative Assessment of the Heterogeneity of PD-L1 Expression in Non-Small-Cell Lung Cancer. JAMA Oncol 2, 46–54.

Mittal, D., Young, A., Stannard, K., Yong, M., Teng, M. W., Allard, B., Stagg, J., and Smyth, M. J. (2014). Antimetastatic effects of blocking PD-1 and the adenosine A2A receptor. Cancer Res 74, 3652–3658.

Morrison, C., Pabla, S., Conroy, J. M., Nesline, M. K., Glenn, S. T., Dressman, D., Papanicolau-Sengos, A., Burgher, B., Andreas, J., Giamo, V., et al. (2018). Predicting response to checkpoint inhibitors in melanoma beyond PD-L1 and mutational burden. J Immunother Cancer 6, 32.

Newman, A. M., Liu, C. L., Green, M. R., Gentles, A. J., Feng, W., Xu, Y., Hoang, C. D., Diehn, M., and Alizadeh, A. A. (2015). Robust enumeration of cell subsets from tissue expression profiles. Nat Methods 12, 453–457.

Newman, A. M., Steen, C. B., Liu, C. L., Gentles, A. J., Chaudhuri, A. A., Scherer, F., Khodadoust, M. S., Esfahani, M. S., Luca, B. A., Steiner, D., et al. (2019). Determining cell type abundance and expression from bulk tissues with digital cytometry. Nat Biotechnol 37, 773–782.

Ni, L., and Dong, C. (2017). New B7 Family Checkpoints in Human Cancers. Mol Cancer Ther 16, 1203–1211.

Nishino, M., Ramaiya, N. H., Hatabu, H., and Hodi, F. S. (2017). Monitoring immune-checkpoint blockade: response evaluation and biomarker development. Nat Rev Clin Oncol 14, 655–668.

O’Connor, J. M., Seidl-Rathkopf, K., Torres, A. Z., You, P., Carson, K. R., Ross, J. S., and Gross, C. P. (2018). Disparities in the Use of Programmed Death 1 Immune Checkpoint Inhibitors. Oncologist 23, 1388–1390.

Ohta, A. (2016). A Metabolic Immune Checkpoint: Adenosine in Tumor Microenvironment. Front Immunol 7, 109.

Ott, P. A., Hodi, F. S., and Robert, C. (2013). CTLA-4 and PD-1/PD-L1 blockade: new immunotherapeutic modalities with durable clinical benefit in melanoma patients. Clin Cancer Res 19, 5300–5309.

Picarda, E., Ohaegbulam, K. C., and Zang, X. (2016). Molecular Pathways: Targeting B7-H3 (CD276) for Human Cancer Immunotherapy. Clin Cancer Res 22, 3425–3431.

Riaz, N., Havel, J. J., Makarov, V., Desrichard, A., Urba, W. J., Sims, J. S., Hodi, F. S., Martin-Algarra, S., Mandal, R., Sharfman, W. H., et al. (2017). Tumor and Microenvironment Evolution during Immunotherapy with Nivolumab. Cell 171, 934–949 e915.

Ribas, A. (2012). Tumor immunotherapy directed at PD-1. N Engl J Med 366, 2517–2519.

Ribas, A., Butterfield, L. H., Glaspy, J. A., and Economou, J. S. (2003). Current developments in cancer vaccines and cellular immunotherapy. J Clin Oncol 21, 2415–2432.

Rizvi, N. A., Hellmann, M. D., Snyder, A., Kvistborg, P., Makarov, V., Havel, J. J., Lee, W., Yuan, J., Wong, P., Ho, T. S., et al. (2015). Cancer immunology. Mutational landscape determines sensitivity to PD-1 blockade in non-small cell lung cancer. Science 348, 124–128.

Robert, C., Schachter, J., Long, G. V., Arance, A., Grob, J. J., Mortier, L., Daud, A., Carlino, M. S., McNeil, C., Lotem, M., et al. (2015). Pembrolizumab versus Ipilimumab in Advanced Melanoma. N Engl J Med 372, 2521–2532.

Robert, C., Thomas, L., Bondarenko, I., O’Day, S., Weber, J., Garbe, C., Lebbe, C., Baurain, J. F., Testori, A., Grob, J. J., et al. (2011). Ipilimumab plus dacarbazine for previously untreated metastatic melanoma. N Engl J Med 364, 2517–2526.

Roh, W., Chen, P. L., Reuben, A., Spencer, C. N., Prieto, P. A., Miller, J. P., Gopalakrishnan, V., Wang, F., Cooper, Z. A., Reddy, S. M., et al. (2017). Integrated molecular analysis of tumor biopsies on sequential CTLA- 4 and PD-1 blockade reveals markers of response and resistance. Sci Transl Med 9.

Rosenberg, J. E., Hoffman-Censits, J., Powles, T., van der Heijden, M. S., Balar, A. V., Necchi, A., Dawson, N., O’Donnell, P. H., Balmanoukian, A., Loriot, Y., et al. (2016). Atezolizumab in patients with locally advanced and metastatic urothelial carcinoma who have progressed following treatment with platinum-based chemotherapy: a single-arm, multicentre, phase 2 trial. Lancet 387, 1909–1920.

Sautes-Fridman, C., Petitprez, F., Calderaro, J., and Fridman, W. H. (2019). Tertiary lymphoid structures in the era of cancer immunotherapy. Nat Rev Cancer 19, 307–325.

Sek, K., Molck, C., Stewart, G. D., Kats, L., Darcy, P. K., and Beavis, P. A. (2018). Targeting Adenosine Receptor Signaling in Cancer Immunotherapy. Int J Mol Sci 19.

Sidaway, P. (2018). Immunotherapy-responsive gastric cancers identified. Nat Rev Clin Oncol 15, 590.

Snyder, A., Makarov, V., Merghoub, T., Yuan, J., Zaretsky, J. M., Desrichard, A., Walsh, L. A., Postow, M. A., Wong, P., Ho, T. S., et al. (2014). Genetic basis for clinical response to CTLA-4 blockade in melanoma. N Engl J Med 371, 2189–2199.

Sokal, R. R., and Rohlf, F. J. (1962). THE COMPARISON OF DENDROGRAMS BY OBJECTIVE METHODS. TAXON 11, 33–40.

Solinas, C., Gu-Trantien, C., and Willard-Gallo, K. (2020). The rationale behind targeting the ICOS-ICOS ligand costimulatory pathway in cancer immunotherapy. ESMO Open 5, e000544.

Sorensen, S. F., Zhou, W., Dolled-Filhart, M., Georgsen, J. B., Wang, Z., Emancipator, K., Wu, D., Busch-Sorensen, M., Meldgaard, P., and Hager, H. (2016). PD-L1 Expression and Survival among Patients with Advanced Non-Small Cell Lung Cancer Treated with Chemotherapy. Transl Oncol 9, 64–69.

Suh, W. K., Gajewska, B. U., Okada, H., Gronski, M. A., Bertram, E. M., Dawicki, W., Duncan, G. S., Bukczynski, J., Plyte, S., Elia, A., et al. (2003). The B7 family member B7-H3 preferentially down-regulates T helper type 1- mediated immune responses. Nat Immunol 4, 899–906.

Tekle, C., Nygren, M. K., Chen, Y. W., Dybsjord, I., Nesland, J. M., Maelandsmo, G. M., and Fodstad, O. (2012). B7-H3 contributes to the metastatic capacity of melanoma cells by modulation of known metastasis-associated genes. Int J Cancer 130, 2282–2290.

Teng, M. W., Ngiow, S. F., Ribas, A., and Smyth, M. J. (2015). Classifying Cancers Based on T-cell Infiltration and PD-L1. Cancer Res 75, 2139–2145.

Tumeh, P. C., Harview, C. L., Yearley, J. H., Shintaku, I. P., Taylor, E. J., Robert, L., Chmielowski, B., Spasic, M., Henry, G., Ciobanu, V., et al. (2014). PD-1 blockade induces responses by inhibiting adaptive immune resistance. Nature 515, 568–571.

Turcu, G., Nedelcu, R. I., Ion, D. A., Brinzea, A., Cioplea, M. D., Jilaveanu, L. B., and Zurac, S. A. (2016). CEACAM1: Expression and Role in Melanocyte Transformation. Dis Markers 2016, 9406319.

Twitty, C. G., Huppert, L. A., and Daud, A. I. (2020). Prognostic Biomarkers for Melanoma Immunotherapy. Curr Oncol Rep 22, 25.

Van Allen, E. M., Miao, D., Schilling, B., Shukla, S. A., Blank, C., Zimmer, L., Sucker, A., Hillen, U., Foppen, M. H. G., Goldinger, S. M., et al. (2015). Genomic correlates of response to CTLA-4 blockade in metastatic melanoma. Science 350, 207–211.

Veenstra, R. G., Flynn, R., Kreymborg, K., McDonald-Hyman, C., Saha, A., Taylor, P. A., Osborn, M. J., Panoskaltsis-Mortari, A., Schmitt-Graeff, A., Lieberknecht, E., et al. (2015). B7-H3 expression in donor T cells and host cells negatively regulates acute graft-versus-host disease lethality. Blood 125, 3335–3346.

Vigano, S., Alatzoglou, D., Irving, M., Menetrier-Caux, C., Caux, C., Romero, P., and Coukos, G. (2019). Targeting Adenosine in Cancer Immunotherapy to Enhance T-Cell Function. Front Immunol 10, 925.

Vilain, R. E., Menzies, A. M., Wilmott, J. S., Kakavand, H., Madore, J., Guminski, A., Liniker, E., Kong, B. Y., Cooper, A. J., Howle, J. R., et al. (2017). Dynamic Changes in PD-L1 Expression and Immune Infiltrates Early During Treatment Predict Response to PD-1 Blockade in Melanoma. Clin Cancer Res 23, 5024–5033.

Wang, L., Le Mercier, I., Putra, J., Chen, W., Liu, J., Schenk, A. D., Nowak, E. C., Suriawinata, A. A., Li, J., and Noelle, R. J. (2014). Disruption of the immune-checkpoint VISTA gene imparts a proinflammatory phenotype with predisposition to the development of autoimmunity. Proc Natl Acad Sci U S A 111, 14846–14851.

Wennhold, K., Shimabukuro-Vornhagen, A., and von Bergwelt-Baildon, M. (2019). B Cell-Based Cancer Immunotherapy. Transfus Med Hemother 46, 36–46.

Wicklein, D., Otto, B., Suling, A., Elies, E., Luers, G., Lange, T., Feldhaus, S., Maar, H., Schroder-Schwarz, J., Brunner, G., et al. (2018). CEACAM1 promotes melanoma metastasis and is involved in the regulation of the EMT associated gene network in melanoma cells. Sci Rep 8, 11893.

Yonesaka, K., Haratani, K., Takamura, S., Sakai, H., Kato, R., Takegawa, N., Takahama, T., Tanaka, K., Hayashi, H., Takeda, M., et al. (2018). B7-H3 Negatively Modulates CTL-Mediated Cancer Immunity. Clin Cancer Res 24, 2653–2664.

Zarour, H. M. (2016). Reversing T-cell Dysfunction and Exhaustion in Cancer. Clin Cancer Res 22, 1856–1864.

